# Clarified an rDNA Gene Unit Pattern with (CTTT)n and (CT)n Microsatellites Aggregation Ahead of and Behind the Gene in Human Genome

**DOI:** 10.64898/2026.03.22.713381

**Authors:** Junxiong Shen, Shiying Tang, Yun Xia, Jieyun Qin, Hanying Xu, Zhongyang Tan

## Abstract

**Background:** Conventional models of human ribosomal DNA (rDNA) array organization have historically depended on transcription-centric boundaries, partitioning the unit into a ∼13 kb rDNA transcription region and a monolithic ∼31 kb intergenic spacer (IGS). While our previous identification of Duplication Segment Units (DSUs) mapped these arrays based on an intuitive analysis of the microsatellite density landscape of the complete reference human genome, our present deep mining of this landscape has revealed a more accurate rDNA Gene Unit Pattern.

**Methods & Results:** In this study, we conducted a deep mining analysis of our previously established microsatellite density landscape of the T2T-CHM13 assembly, focusing specifically on nucleolar organizing regions (NORs). We suggest a more accurate rDNA Gene Unit Pattern containing a (CTTT)n microsatellite aggregation ahead of the rDNA gene and a (CT)n microsatellite aggregation behind the gene, rather than a pattern featuring an IGS region inserted between two rDNA genes.

**Conclusions:** A correct rDNA gene pattern of the human genome probably includes a (CTTT)n microsatellite aggregation ahead of the gene and a (CT)n microsatellite aggregation behind it, which possibly constitute cis- and trans-regulating regions; the (CTTT)n and (CT)n microsatellite aggregations may provide two different local stable DNA structures for regulatory protein binding.

## 1. Introduction

The organization and stability of the human genome are profoundly influenced by the structure and dynamics of ribosomal DNA (rDNA) arrays, which rank among the most transcriptionally active and repetitive loci in the genome. Located in nucleolar organizing regions across the short arms of five acrocentric chromosomes, these loci are composed of hundreds of tandem repeats (Henderson et al., 1972; Sylvester et al., 1986). The active transcription of rRNA genes forms the structural and functional core of the nucleolus, a multifunctional nuclear body central to cell cycle progression, cellular homeostasis, and genome integrity (Pederson, 2011; Grummt, 2013). However, the highly repetitive nature of rDNA, particularly within the intergenic spacer (IGS), has historically posed significant challenges for genome assembly and bioinformatic analysis. This technical barrier resulted in fragmented representations in standard references, hindering comprehensive studies of sequence variation, recombination, and the functional roles of repetitive elements (Treangen & Salzberg, 2012). Consequently, early characterizations of the rDNA repeat unit relied on idealized consensus models or context specific sequences from isolated clones, providing a refined yet fundamentally incomplete picture of the locus.

Overcoming these barriers is critical because the IGS is far from a mere structural void. It is a known hotspot for recombination and structural variation, harboring a diverse array of repetitive elements including microsatellites, minisatellites, and transposable elements (Wahls et al., 1990; Stults et al., 2008; Bendich, n.d., 2023). Beyond its role in driving rapid genomic evolution and potential instability (Smirnov et al., 2021), the IGS contains essential regulatory sequences, protein binding sites, and noncoding RNAs that actively influence rDNA transcription, nucleolar organization, and cellular stress responses (Vacík et al., 2019). Among these diverse components, microsatellites or short tandem repeats have emerged as uniquely potent functional elements. Rather than acting as neutral genomic filler, these sequences directly modulate gene regulation, chromatin folding, and three dimensional architectural organization (Kumar et al., 2013). Furthermore, their highly mutable nature makes them powerful drivers of sequence heterogeneity (Kashi & King, 2006). This functional versatility strongly suggests that the microsatellite distribution within the IGS serves as a critical architectural code necessary for fine tuning rDNA activity and maintaining genomic stability.

The completion of the telomere to telomere T2T CHM13 human genome assembly fundamentally transformed this research landscape by providing the first contiguous, gapless reference for these highly complex regions (Antonarakis, 2022). This milestone enabled the precise mapping of the microsatellite landscape at single nucleotide resolution. Yet, despite this comprehensive reference, much of the subsequent research has remained conceptually fragmented, frequently isolating specific sequence elements or localized regions rather than adopting a whole genome perspective. Because genomic regulation operates through continuous spatial networks, adopting a comprehensive structural perspective is essential to systematically compare repeat characteristics and distinguish functionally conserved motifs from mere accumulative variations across diverse genomic contexts.

Applying this global perspective, our previous mapping of the whole genome microsatellite distribution uncovered a striking architectural feature: pronounced palisading clusters of CT repeats specifically localized to the short arms of all five acrocentric chromosomes (Xia et al., 2024). This genomic pattern indicates that despite their inherent length variability, the sheer accumulation of these repeats may serve a profound structural purpose. Rather than relying on a precise sequence length, the clustering itself could facilitate the formation of specific higher-order configurations, such as distinct double-stranded structures. Consequently, these formations can act as physical boundaries that partition functional genomic domains to regulate regional gene expression. However, while our initial mapping successfully identified this genome-wide microsatellite accumulation, the precise internal topology of the rDNA arrays and the exact spatial relationship between the rRNA coding sequences and their surrounding IGS regulatory domains required a more targeted investigation.

In this study, we employ an in silico approach to comprehensively delineate the micro-architecture of microsatellite clustering specifically within the rDNA arrays of the T2T-CHM13 assembly. By quantitatively mapping the distribution of these motifs across all rDNA repeat units, we identified two distinct types of microsatellite clustering peaks that stably and preferentially localize within the intergenic spacer (IGS). Rather than randomly accumulating, these specific aggregations display a strict spatial orientation relative to the coding regions. The precise localization of these distinct microsatellite clusters challenges the traditional view of the IGS as a monolithic spacer, suggesting they potentially serve as critical architectural boundaries of the rDNA repeating unit. Ultimately, this high-resolution sequence analysis establishes a robust one-dimensional coordinate system, providing a precise sequence foundation to guide future experimental investigations into the 3D regulatory architecture of these fundamental genomic domains.

## 2. Materials and methods

### 2.1. Deep mining the microsatellite density landscape

We performed a deep mining of the microsatellite density landscape based on our previous work (Xia et al., 2024). This analysis was conducted across all chromosomes, with a particular focus on the Nucleolar Organizing Regions (NORs).

### 2.2. Genome sequences

The complete set of 24 chromosome sequences from the human telomere-to-telomere (T2T) reference genome CHM13v2.0 was obtained from GenBank. NORs coordinates were defined using the CenSat annotation track available through the UCSC Genome Browse.

### 2.3. Detection of a Conserved (CTTT)n-Rich Regulatory Head at the Physical Start of rDNA Arrays

To delineate the true physical origin of human ribosomal DNA (rDNA) repeating units, we first defined the boundaries of NORs on the five acrocentric chromosomes (13, 14, 15, 21, and 22) using the CenSat annotation track from the UCSC Genome Browser. For each chromosome, we extracted a reference window extending from the annotated NOR start coordinate to the transcription start site (TSS) of the first 45S rDNA gene and further expanded it by 10 kb upstream, yielding a ∼14 kb region that fully encompasses the transition between pericentromeric heterochromatin and the rDNA array.

This reference window served as the search space for identifying the conserved “regulatory head.” For each of the 219 rDNA repeat units, we extracted the terminal ∼5 kb of its intergenic spacer (IGS)—corresponding to the segment immediately upstream of the 45S coding sequence—and aligned it against the chromosome-matched reference window using Clustal Omega (v1.2.4) with default parameters for nucleotide sequences. The subregion within the reference window showing the highest local sequence similarity to the IGS terminus was designated as the candidate regulatory head for that unit.

To assess the structural features of these candidates, we integrated the alignment results with the microsatellite density landscape. We examined whether each candidate overlapped a high- or medium-density microsatellite peak of the (CTTT)n motif [H/MMDP-(CTTT)n], a common feature of canonical rDNA architecture, but this was used only for descriptive purposes. The definition of an rDNA genomic unit (rDNA-GU) was based solely on significant sequence conservation between the candidate region and the IGS terminus. All 219 rDNA units showed sufficient local sequence similarity and were retained for subsequent structural modeling and functional annotation.

Pairwise sequence identity values derived from the Clustal Omega alignments are summarized in Table 1, and the conservation profile across all identified regulatory heads is visualized in the heatmap in Supplementary Figure S3.

**Table 1.**
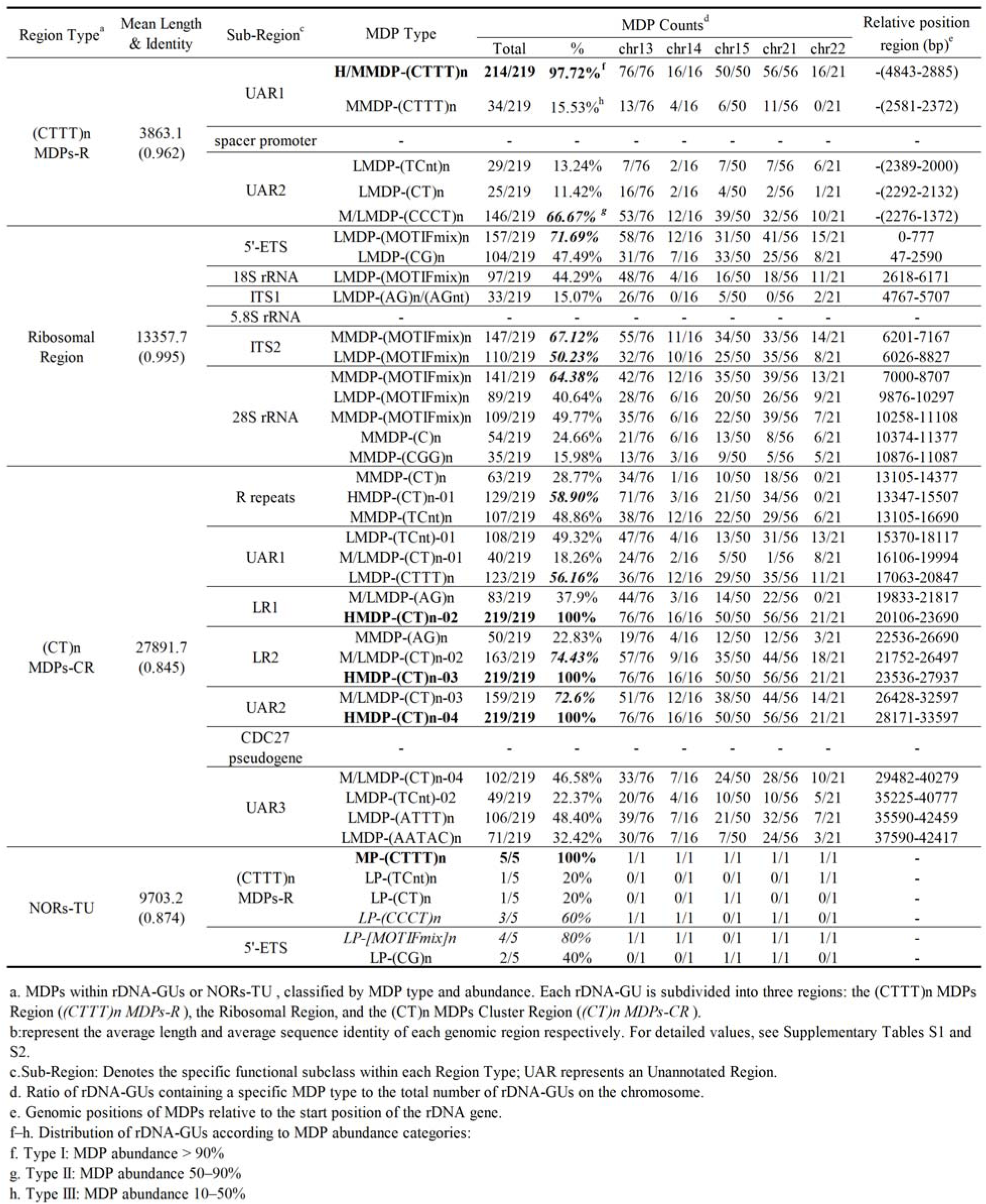
A Microsatellite Density Peak (MDP) Distribution Pattern in rDNA-GUs and NORs-TUs.

### 2.4. Characterization of NOR architecture using microsatellite density landscapes and sequence alignment

Following the identification of the (CTTT)n-rich regulatory head at the physical start of each rDNA array, we systematically delineated the full structural architecture of NORs in the T2T-CHM13 genome.

Using the microsatellite density landscape maps generated in our prior study (Xia et al., 2024) as a structural scaffold, we annotated all rDNA-containing segments on chromosomes 13, 14, 15, 21, and 22.

For each of the 219 validated rDNA genomic units (rDNA-GUs), we defined three consecutive structural domains:

i. an upstream (CTTT)n microsatellite density peak region (MDP-R), corresponding to the conserved regulatory head identified in Section 2.3;
ii. the core rDNA transcriptional unit, encompassing the 45S pre-rRNA coding sequence (including 18S, 5.8S, and 28S rRNA genes along with internal transcribed spacers);
iii. a downstream (CT)n MDP cluster region (MDP-CR), characterized by a dense array of (CT)n microsatellite peaks.

To assess structural conservation across the array, we performed multiple sequence alignments of all rDNA-GUs using Clustal Omega (v1.2.4) with default parameters for nucleotide sequences.

### 2.5. Identification and Structural Characterization of the NORs Terminal Unit (NORs-TU)

Finally, based on the NOR boundaries defined by the CenSat annotation track from the UCSC Genome Browser, we identified a single NORs terminal unit (NORs-TU) within the distal segment of each acrocentric chromosome’s short arm, specifically in the region immediately downstream of the last complete rDNA-GU.

In total, we annotated 219 full-length rDNA-GUs and 5 NORs-TUs across the five acrocentric chromosomes: All annotated coordinates, domain boundaries, and motif classifications are provided in Supplementary Tables S1.01–S1.06.

### 2.6. Construction of a MDP Distribution Pattern for rDNA-GUs

We developed a custom Python-based tool, Microsatellite Unit Profiler v1.0 (available at: https://github.com/zhongyangtan/MUP.git), to automate the profiling of microsatellite density peaks (MDPs) within individual rDNA-GUs. Based on the proportion of the rDNA-GU occupied by MDPs, units were classified into three categories:

Type I: MDP abundance > 90%
Type II: MDP abundance 50–90%
Type III: MDP abundance 10–50%

Detailed classifications and unit-level statistics are provided in Supplementary Tables S3.01–S3.03.

### 2.7. Comparison of MDP-based rDNA-GU classification with historical IGS and DSU models

Our MDP distribution patterns were compared with the IGS regions described in prior literature as well as with the DSUs defined in our previous study(Xia et al., 2024). Additionally, comparisons were made against the reference human ribosomal RNA repeat unit (GenBank accession: KY962518), which includes annotated features such as the Spacer Promoter, R repeats, LR1, LR2, and the CDC27 pseudogene. Sequence alignments supporting these comparisons were generated using Clustal Omega and are documented in Supplementary Tables S2.04–S2.06.

### 2.8. Generation of composite and individual (CT)n/(CTTT)n landscape maps

Following the methodology established in Xia et al. (2024), we constructed genome-wide Microsatellite density landscape maps for the T2T-CHM13 assembly, focusing on three track types: (i) regions containing both (CT)n and (CTTT)n MDPs, (ii) regions with only (CT)n MDPs, and (iii) regions with only (CTTT)n MDPs. These tracks have been deposited in the UCSC Genome Browser and are publicly accessible via the following link: [https://genome.ucsc.edu/s/zhongyangtan/CHM13v2.1].

### 2.9. Definition and Statistical Characterization of (CT/CTTT)n MDP-Depleted Genomic Intervals

We developed MDP-ZoneScanner, a Python-based tool for genome-wide detection of regions devoid of (CT)n or (CTTT)n microsatellite density peaks (MDPs) (available at: https://github.com/zhongyangtan/MDP-ZoneScanner.git). Applying this tool to the T2T-CHM13 assembly, we identified and classified MDP-depleted intervals into two categories:

- Common Zero (CT/CTTT)n MDP Regions (Co-0 (CT/CTTT)n MDP-Rs): intervals >3 Mb with no detectable (CT)n or (CTTT)n MDPs;
- Common Zero (CT/CTTT)n MDP Big Regions (Co-0 (CT/CTTT)n MDP-BRs): intervals >10 Mb entirely lacking such MDPs.

Genomic maps of these regions are provided in Supplementary Figures S5.01–S5.03.

### 2.10. Correlation mapping between HMDP-(CT)n/HMDP-(CTTT)n and gene annotations

We developed HMDP-GeneMapper, a Python-based pipeline to systematically assess the spatial association between high-density microsatellite peaks [HMDP-(CT)n and HMDP-(CTTT)n] and gene annotations from the UCSC CAT/Liftoff Genes track (available at: https://github.com/zhongyangtan/HMDP-GeneMapper.git). The tool catalogs all genes that either overlap an HMDP region or lie within ±100 kb of its boundaries, and assigns them to functional categories for downstream analysis.

## 3. Result

Prior research established a specialized framework for parsing ribosomal DNA (rDNA) arrays using microsatellite density peaks (MDPs), premised on the observation that a systematic enrichment of high-density microsatellite peaks (HMDPs) within non-coding regions, 89.25% in intergenic spacers (IGS) and 10.74% in introns (Xia et al., 2024). This pattern suggests that microsatellite clusters serve as fundamental scaffolds for demarcating genomic structural units. Specifically, we identified HML(CT)n cluster regions, which are comprised of HMDPs with (CT)n motifs arranged in a palisade fashion alongside MMDPs and LMDPs, and which define the tandem Duplication Segment Unit (DSU). Across the five human acrocentric chromosomes, we identified 219 DSUs (ranging from 43,000 to 47,000 bp), where [CT]n HMDP groups precisely separate individual 45S rDNA gene clusters.

However, while the DSU model provided a high-resolution map of the repeat units, it largely deferred to the conventional partition of rDNA, which divides the unit into a ∼13 kb transcription region and a ∼31 kb IGS (Wellauer & Dawid, 1979). Consequently, the initial DSU definition treated the IGS as a monolithic spacer and placed the physical boundary at the Transcription Start Site (TSS). In this study, a high-resolution re-examination of the microsatellite landscape reveals that this partition is structurally inaccurate. We demonstrate that the IGS is not a uniform spacer but comprises distinct upstream and downstream regulatory domains, necessitating a revision of the DSU boundary to reflect the true physical organization of the rDNA array.

### 3.1. Identification of an Overlooked “Regulatory Head” at the rDNA Array Starts

A high-resolution analysis of the microsatellite landscape across the nucleolar organizer regions (NORs) of the T2T-CHM13 genome revealed a conserved structural element absent from all previous models. We identified a distinct H/MMDP−(CTTT)n peak at the absolute physical commencement of the NOR on all five acrocentric chromosomes (Figure 1B shows chromosome 14; other chromosomes are shown in Supplementary Figure 1), located approximately 4,000 bp upstream of the first rDNA gene. Moreover, this upstream (CTTT)n peak presents in 97.72% of all rDNA genes (Table 1), except five instances on chromosome 21 (Supplementary Table S3).

**Figure 1.**
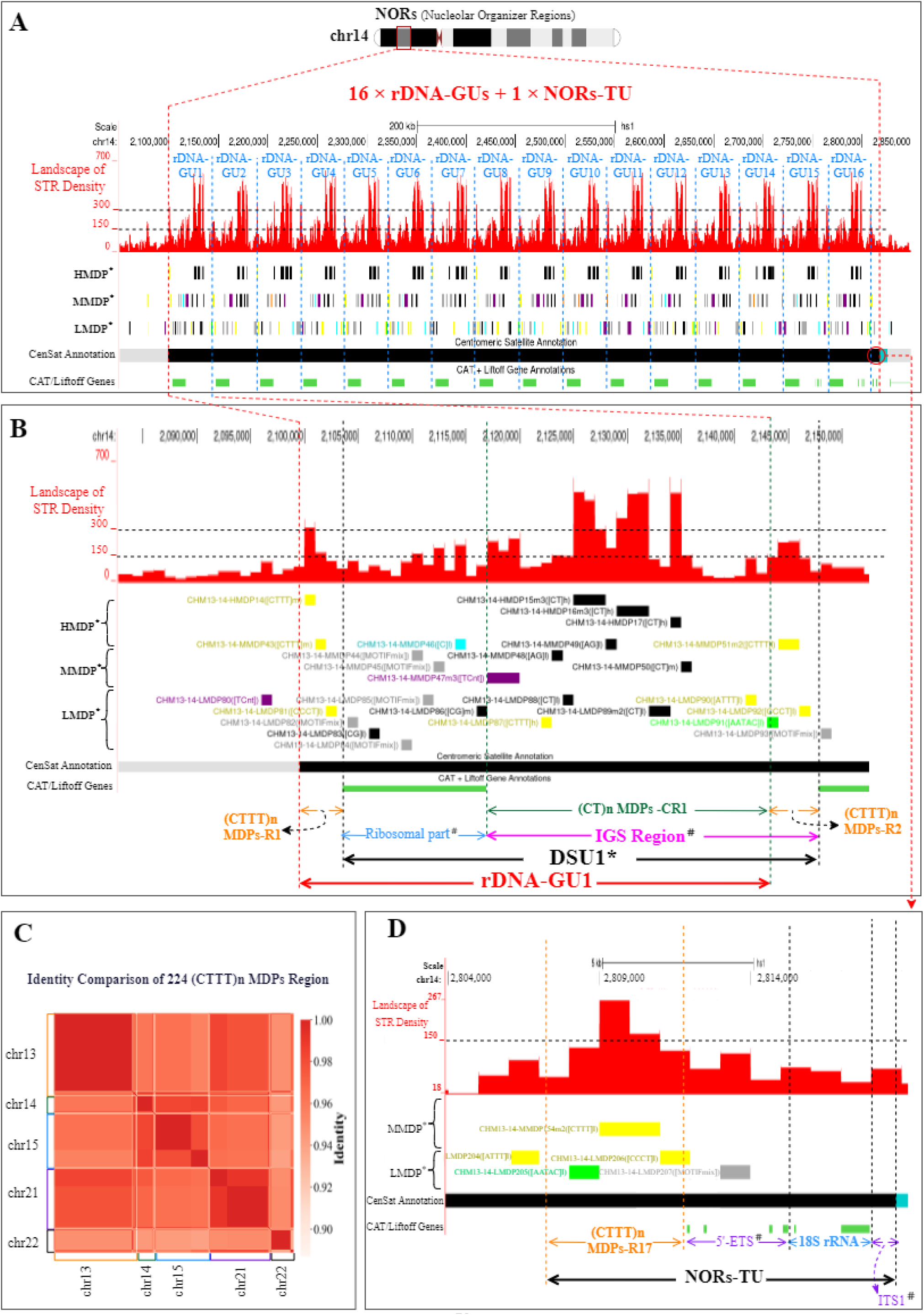
The human nucleolar organizer regions (NORs) are composed of multiple rDNA genomic units (rDNA-GUs) and a terminal unit (NORs-TU). **A.** Overview of the NORs on chromosome 14, showing 16 rDNA-GUs and one NORs-TU, annotated with microsatellite density landscape and microsatellite density peaks (HMDPs, MMDPs, LMDPs). rDNA gene clusters and NORs positions are shown using UCSC tracks (CAT/Liftoff for rDNA; CenSat Annotation for NORs). B. Detailed view of a single rDNA-GU, revealing its tripartite structure: (CTTT)n MDP-R, rDNA transcriptional unit, and downstream (CT)n MDP-CR. C. Pairwise sequence identity heatmap of all (CTTT)n MDP-R regions across five acrocentric chromosomes (n=224), demonstrating high conservation. D. Microsatellite density profile of the NORs-TU, showing retention of (CTTT)n MDP-R but absence of downstream (CT)n MDP-CR and variable truncation of the rDNA unit. Legend: Structural or organizational features defined in Xia et al. (2024) HMDP: High microsatellite density peak MMDP: Medium microsatellite density peak LMDP: Low microsatellite density peak DSU: Duplication segment unit, comprising a rDNA unit and an IGS Functional elements within the rDNA transcriptional unit: 5′-ETS: 5′ external transcribed spacer 18S rRNA: 18S ribosomal RNA gene ITS1: Internal transcribed spacer 1 IGS: Intergenic spacer (non-transcribed region separating rDNA repeats)

Sequence alignment revealed that this ∼4,000 bp segment at the NOR start shares over 90% similarity with the terminal ∼4,000 bp of every DSU (i.e., the end of the conventional IGS) (Figure 1C; detailed heatmap in Supplementary Figure 3). This significant homology demonstrates that the rDNA array does not physically initiate with the ribosomal gene itself, but rather with a ∼4,000 bp (CTTT)n-rich segment, a "regulatory head" that sets the structural stage for the subsequent array.

### 3.2. Revising the rDNA Unit Model: Beyond Transcription-Centric Boundaries

This discovery underscores the fundamental limitations of prevailing rDNA structural models. Historically, the definition of the rDNA unit was constrained by fragmented, cloned sequences, engendering a "transcription-centric" perspective. This paradigm rested on a coarse binary distinction between functional coding domains and inert non-coding regions, thereby obscuring the structural complexity of the intergenic intervals, particularly the architectural significance of repetitive elements. Consequently, this approach erroneously designated the Transcription Start Site (TSS) as the unit’s physical origin. While the T2T-CHM13 genome now enables the delineation of NOR boundaries via sequence similarity (Nurk et al., 2022), it lacks the requisite resolution to decode the internal structural logic.

Our microsatellite density landscape provides this missing resolution. Our previous DSU model, though an improvement, focused primarily on internal repeat patterns and failed to account for the boundary signals that define the true origin of the unit. The current findings allow us to refine this concept:

1. Reconceptualization of the IGS: The conventional IGS should no longer be viewed as a monolithic "spacer." Instead, it is partitioned into two distinct subdomains: one located downstream of the gene cluster and another positioned upstream.
2. Structural Revision: The traditional "gene-spacer" dichotomy oversimplifies the rDNA array as a series of functional genes separated by non-functional gaps. Our model supports a more nuanced architecture where the (CTTT)n-rich "head" acts as a conserved upstream regulatory reservoir.

By integrating these boundary markers with internal MDP patterns, we provide a structural foundation that more accurately reflects the physical and regulatory reality of the NOR. This revised model suggests that the coordinated transcription of rDNA is governed by a bidirectional regulatory logic, with the "head" segment playing a potentially crucial role in organizing the chromatin landscape of the array.

### 3.3. The rDNA Gene Unit (rDNA-GU): A Tripartite Structural Model

To identify the most stable structural components within this new framework, we generated a consensus pattern across all 219 rDNA-GUs in the T2T-CHM13 genome (Figure 2). We filtered for peaks present in at least 10% of the rDNA-GUs to ensure robustness against microsatellite instability. This systematic analysis revealed a highly conserved, tripartite architecture underlying the ribosomal array.

**Figure 2.**
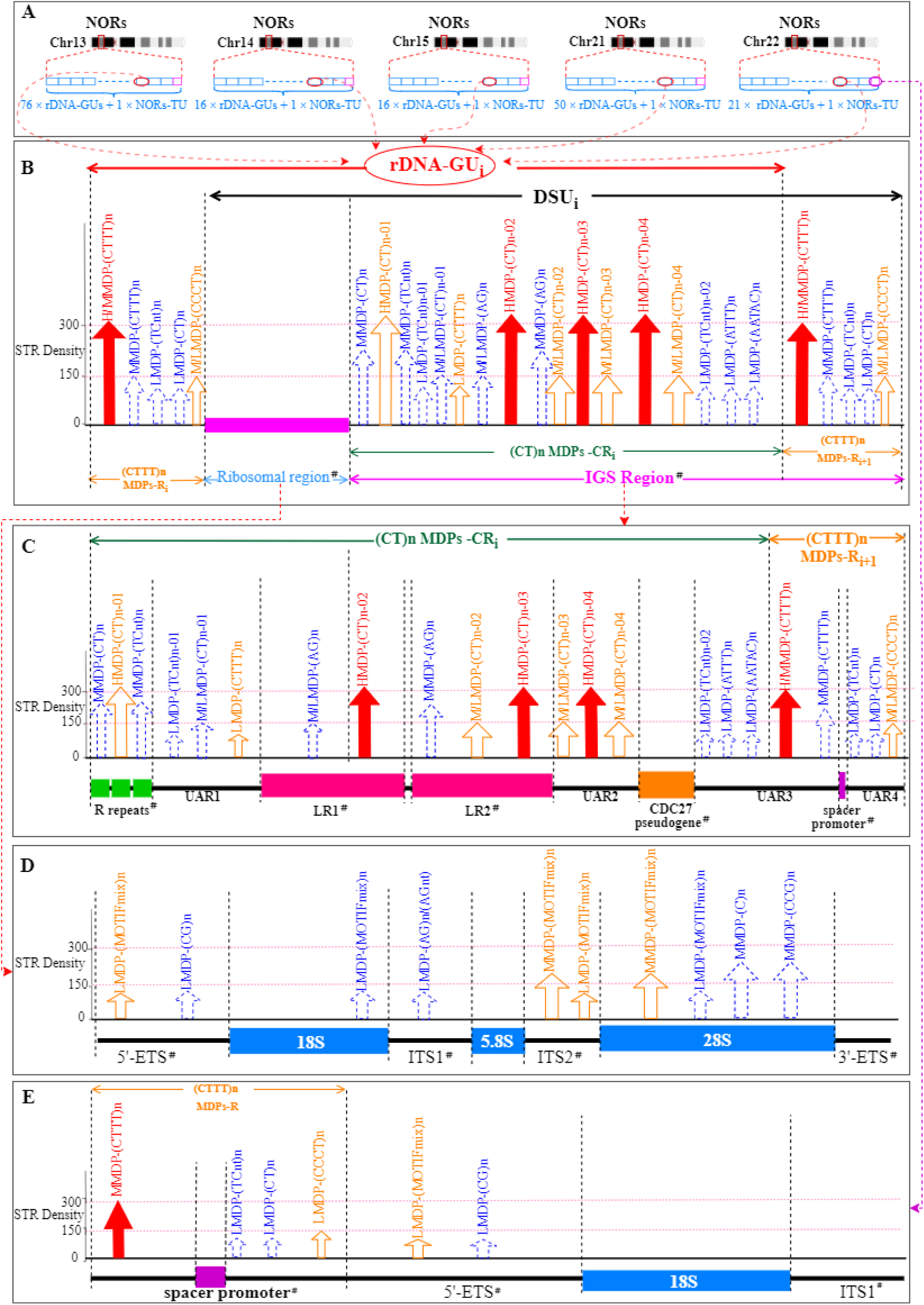
A Microsatellite Density Peaks (MDPs) Pattern in rDNAs-GU and NORs-TU. **A.** Schematic representation of nucleolar organizer regions (NORs) on human acrocentric chromosomes 13, 14, 15, 21, and 22 in the T2T-CHM13 assembly, showing variable numbers of rDNA genomic units (rDNA-GUs) followed by a single terminal unit (NORs-TU).**B.** Schematic of the rDNA genomic units (rDNA-GUs) showing the distribution of MDPs across the full repeating unit. **C.** Schematic of the intergenic spacer (IGS) showing the distribution of MDPs across the non-coding region. **D.** Schematic of the ribosomal part showing the distribution of MDPs across the rRNA gene unit.**E.** Schematic of the NOR terminal units (NORs-TUs) showing the distribution of MDPs across the truncated terminal segment. Legend: 5′-ETS: 5′ external transcribed spacer 18S rRNA: 18S ribosomal RNA gene ITS1: internal transcribed spacer 1 5.8S rRNA: 5.8S ribosomal RNA gene ITS2: internal transcribed spacer 2 28S rRNA: 28S ribosomal RNA gene 3′-ETS: 3′ external transcribed spacer (after 28S) R repeats: regulatory repeats in the IGS LR1/LR2: long repeat ½ CDC27 pseudogene: CDC27 processed pseudogene (IGS-embedded) Spacer promoter: IGS promoter driving non-coding RNA transcription and rDNA silencing

Based on this distinct landscape, we propose the rDNA Gene Unit (rDNA-GU) model to more accurately capture the complete physical and functional architecture of the repeating unit. Unlike the traditional gene-spacer dichotomy, the rDNA-GU is defined by its structural landmarks and comprises three consecutive domains:

1. The Upstream (CTTT)n MDPs-R: Initiated by a H/MMDP−(CTTT)n peak, this region spans an average of 3,864 bp. It exhibits remarkable sequence conservation across the array (mean identity = 0.962) and serves as the structural "head" of the unit. The H/MMDP-(CTTT)n peak at the start of unit serves as the second most consistent structural landmark in our study, appearing in 97.72% of all units (Table 1). Following this anchor, the region encompasses several additional peaks that correspond to the known enhancer repeat elements. While these intermediate peaks exhibit greater positional variability than the initial (CTTT)n landmark, they collectively maintain a coherent compositional gradient. Notably, as the sequence approaches the 18S gene boundary, we observe a characteristic shift in nucleotide composition: a gradual decrease in Thymine accompanied by an increase in Cytosine content. This gradient culminates in a conserved M/LMDP-(CCCT)n peak (present in 66.67% of units), located just upstream of the transcription start site (TSS), marking the precise transition into the gene-coding region.
2. The Core Transcription Region: Comprising the 18S, 5.8S, and 28S rRNA gene clusters. As anticipated, the gene-coding regions showed a scarcity of repeat enrichment. Merely four M/LMDPs located within the 5’ETS, ITS2, and 28S regions exceeded 50% occurrence frequency, with no dominant motif type clustering in these areas (Figure 2D).
3. The Downstream (CT)n MDP-CR: Spanning approximately 27,892 bp, this region is characterized by the clustering of (CT)n MDPs. While longer and more variable than the (CTTT)n MDPs-R, it maintains a high mean sequence identity of 0.845.

The downstream regulatory region is characterized by CT-rich sequences and anchored by a core framework of three highly conserved HMDP-(CT)n peaks with 100% occurrence frequency (Figure 2C). Beyond this invariant core, several other microsatellite peaks show occurrence frequencies above 50% and define key architectural landmarks. Notably, a single (CTTT)n LMDP precedes the first long repeat (LR1), and a HMDP-(CT)n peak (63.47% frequency) marks the R repeat. All other conserved peaks above this frequency threshold reside within the Long Repeat regions. The long repeats themselves exhibit comparable peak compositions: LR1 is characterized by M/LMDP-(AG)n and HMDP-(CT)n peaks, while LR2 displays a similar profile reinforced by an additional M/LMDP-(CT)n peak.

Under this refined framework, the human NOR can be parsed into a contiguous series of complete rDNA-GUs, followed by a single NOR Terminal Unit (NOR-TU) at the proximal end. We identified 76 rDNA-GUs on chromosome 13, 50 on chromosome 21, and 21 on chromosome 22, with chromosomes 14 and 15 each harboring 16 units. The NOR-TU represents a truncated repeat; for instance, the terminal unit on chromosome 14 contains only the 5’ETS, 18S rRNA, and ITS1 regions (Figure 1D; other chromosomes are shown in Supplementary Figure 2). The existence of this consistent terminal truncation suggests a programmed structural boundary for the array, though its precise mechanistic origin remains to be fully elucidated.

### 3.4. Genomic Specificity and Architectural Significance of (CT)n and (CTTT)n

To determine whether the predominance of (CT)n peak clusters and (CTTT)n solitary peaks is a feature specific to the NOR or a widespread genomic one, we mapped the global distribution of microsatellite density peaks (MDPs) across the entire T2T-CHM13 genome. This analysis revealed a distinct, chromosome-specific landscape (Figure 3A, Supplementary Figure4). Generally, centromeric regions across all chromosomes are depleted of microsatellite peaks, likely due to the dominance of long-range repeats such as α-satellite (Altemose et al., 2022). On the five acrocentric chromosomes, however, the short arms exhibit a unique, tripartite zonal architecture (Figure 3B). The distal region (closest to the telomere) is characterized predominantly by Low-Density Peaks (LMDPs). This transitions into the central NOR region, which is defined by a highly regular, “periodic” arrangement of High-Density Peaks (HMDPs). Following the NOR, the proximal region (approaching the centromere) initially displays a segment of irregular mixed density peaks before tapering off into a final low-density zone similar to the distal end.

**Figure 3.**
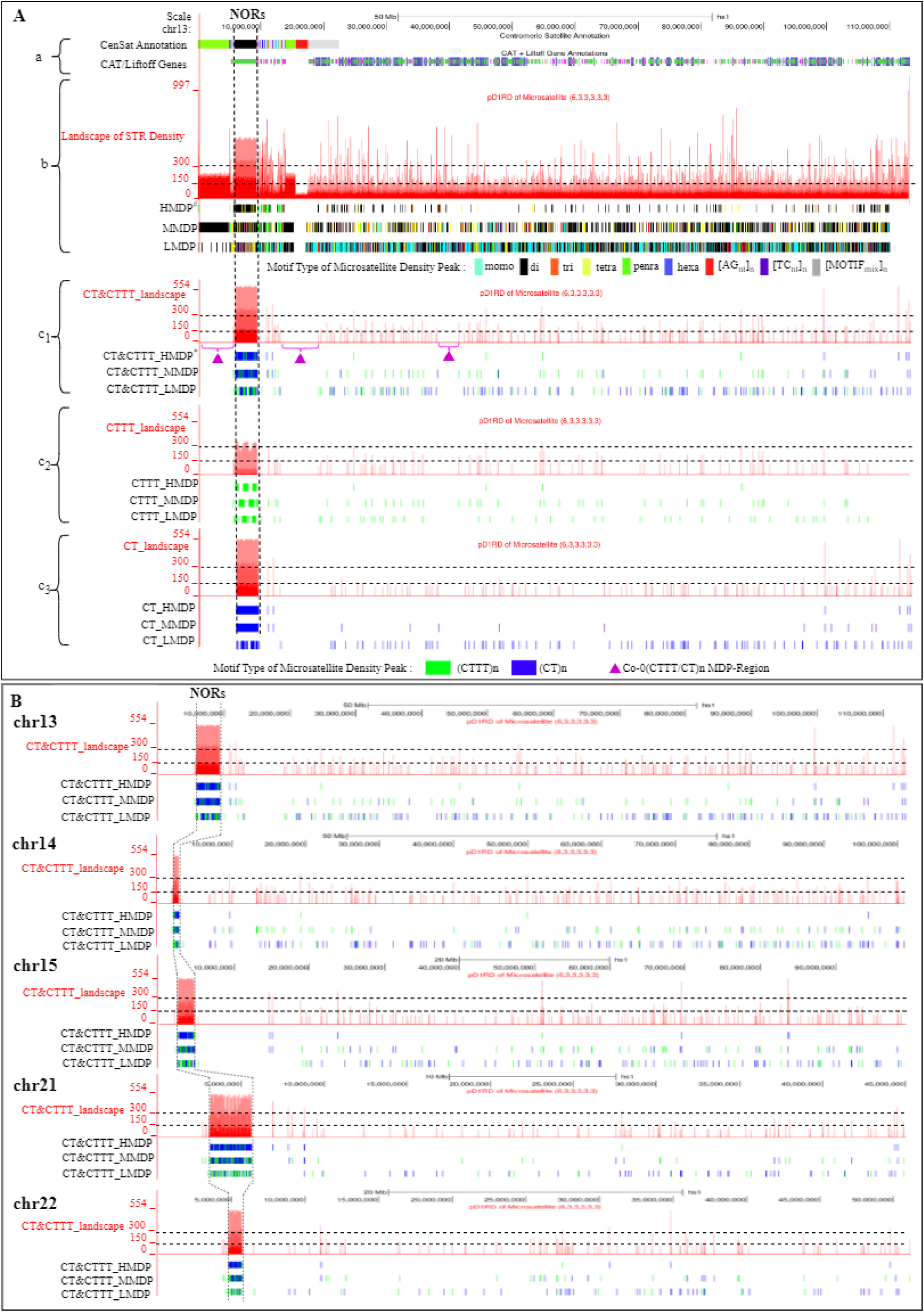
[CT]n and [CTTT]n Microsatellite Density Peaks Show Higher-Density Clustering in Nucleolar Organizer Regions Than in Other Genomic Regions. **(A)** Detailed views of chromosome 13 tracks: **a.** UCSC CAT/Liftoff gene annotations and CenSat repeat annotations, displayed in yellow and orange, respectively. **b.** Overall Microsatellite density profile across chromosome 1, color-coded according to repeat motif classes: mononucleotide (cyan), dinucleotide (black), trinucleotide (orange), tetranucleotide (yellow), pentanucleotide (light green), hexanucleotide (blue), [AGn]n (red), [TCn]n (purple), and mixed motifs ([MOTIFmix]n, gray).**c.**Overview of [CT]n and [CTTT]n MDP distributions. (c_1_) Combined view with type-specific coloring ([CT]n: blue; [CTTT]n: green). (c_2_) Isolated distribution of [CTTT]n MDPs (green). (c_3_) Isolated distribution of [CT]n MDPs (blue). **(B)** Microsatellite density landscapes of [CT]n (blue) and [CTTT]n (green) across chromosomes 13, 14, 15, 21, and 22, annotated with High (HMDPs), Middle (MMDPs), and Low (LMDPs) microsatellite density peaks.

Within this structured landscape, (CT)n and (CTTT)n motifs emerge as the specific drivers of the stable, high-density clustering observed exclusively in the NORs. The quantitative disparity between the NORs and the rest of the genome is striking (Supplementary Table 2). Globally, we identified 905 (CT)n HMDPs, of which an astounding 87.1% are sequestered within the NORs. Similarly, 54.9% of the 215 global (CTTT)n HMDPs reside within these regions. When normalized by sequence length, the spatial density of (CT)n and (CTTT)n HMDPs in the NORs is 2,093-fold and 378-fold higher, respectively, than in the non-NOR genome. This overwhelming concentration confirms that these motifs are not merely repetitive noise but are fundamental architectural components specific to the nucleolus organizer regions.

The exclusivity of these motifs to the rDNA array is further underscored by the identification of extensive genomic intervals devoid of such signals. We cataloged Common Zero (CT/CTTT)n Microsatellite Density

Peak Regions (Co-0 (CT/CTTT)n MDP-Rs), which are defined as genomic intervals exceeding 3 Mb that completely lack detectable peaks for these repeats (Supplementary Table S5). Remarkably, four massive Co-0 (CT/CTTT)n MDP-BRs (Big Regions >10 Mb) were identified on chromosomes 1, 9, 16, and Y. Even in the microsatellite-rich regions of non-acrocentric autosomes and sex chromosomes, these specific motifs fail to form the dense clusters characteristic of the NOR. For instance, on chromosome 13, aside from the distal end of short arm, we identified a ∼4 Mb void in the centromeric region and a 379 kb segment in the distal short arm that are entirely free of (CT)n and (CTTT)n clusters.

Finally, to assess whether these clusters serve broader regulatory functions beyond the nucleolus, we analyzed all protein-coding genes within a 10 Mb window of (CT)n and (CTTT)n HMDP sites (Supplementary Table S6). Across the genome, 120 out of 215 (CTTT)n HMDPs are located in the immediate vicinity of rDNA-related clusters (Figure 4). In contrast, the few high-density peaks found outside the acrocentric short arms showed no association with any shared functional pathways or gene families. These rare occurrences appear stochastic, occurring near randomly distributed genes without forming a consistent regulatory pattern. This extreme specificity suggests that the (CT)n and (CTTT)n signatures are unique architectural hallmarks of the human rDNA array, likely evolved to maintain the unique chromatin environment or meet the intense transcriptional demands of the nucleolus.

**Figure 4.**
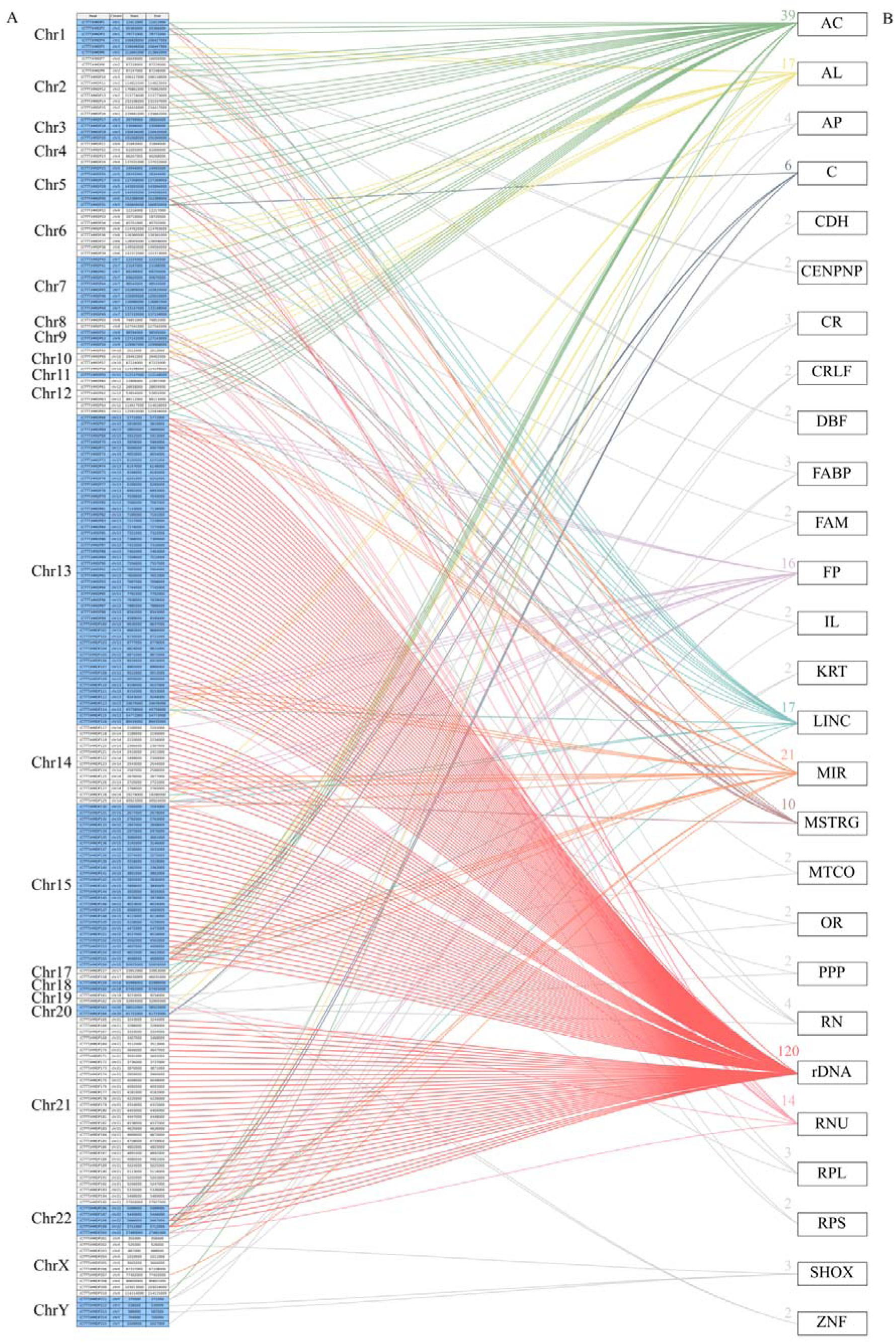
Preferential Enrichment of [CT]n and [CTTT]n HMDPs in Ribosomal DNA (rDNA) Loci Across the Genome. (A) Genome-wide chromosomal distribution of [CTTT]n HMDPs. (B) Mapping of [CTTT]n HMDPs to genes within ±0.1 Mb flanking regions (annotated from UCSC CAT/Liftoff), highlighting a strong association with rDNA.

**Table 2.** Enrichment of [CT]n and [CTTT]n Microsatellite Density Peaks in Nucleolar Organizer Regions(NORs) Relative to the Rest of the T2T-CHM13 Genome.

| MDP Type | Total genome |  |  | NORs |  |  |  | Non-NORs |  |  |  | NORs /<br>Non-NORs<br>Ratio<sup>b</sup> |
| --- | --- | --- | --- | --- | --- | --- | --- | --- | --- | --- | --- | --- |
|  | Length | count | MDP<br>density<sup>a</sup> | Length | count | % | MDP<br>density | Length | count | % | MDP<br>density |  |
| (CTTT)<sub>n</sub>-HMDP |  | 215 | **0.07** |  | 118 | 54.9% | **11.8** |  | 97 | 45.1% | **0.03** | **378** |
| (CTTT)<sub>n</sub>-MMDP |  | 1642 | **0.53** |  | 141 | 8.6% | **14.1** |  | 1501 | 91.4% | **0.48** | **29** |
| (CTTT)<sub>n</sub>-LMDP | 3117 | 1198 | **0.38** | 10 | 138 | 11.5% | **13.8** | 3107 | 1060 | 88.5% | **0.34** | **40** |
| (CT)<sub>n</sub>-HMDP | Mb | 905 | **0.29** | MB | 788 | 87.1% | **78.8** | Mb | 117 | 12.9% | **0.04** | **2093** |
| (CT)<sub>n</sub>-MMDP |  | 1193 | **0.38** |  | 481 | 40.3% | **48.1** |  | 712 | 59.7% | **0.23** | **210** |
| (CT)<sub>n</sub>-LMDP |  | 3569 | **1.15** |  | 122 | 3.4% | **12.2** |  | 3447 | 96.6% | **1.11** | **11** |

a: Microsatellite Density Peak (MDP) density is calculated as the number of MDP per megabase (count/Mb). Total genome size $\approx 3117$ Mb, Nucleolar Organizer Regions (NORs) size $\approx 10$ Mb, non-Nucleolar Organizer Regions(non-NORs) size $\approx 3107$ Mb.
b: Ratio of MDP density in NORs to that in non-NORs (NORs MDP density / non-NORs MDP density).

## 4. Discussion

### 4.1. The Upstream (CTTT)n-Rich "Head" Operates as an Extended Cis-Regulatory Region

A central conclusion of our rDNA-GU model is the reclassification of the ∼4,000 bp (CTTT)n-rich segment, which has been traditionally viewed as the distal tail of the Intergenic Spacer (IGS), into the physical and functional "head" of the rDNA repeat unit. This architectural realignment resolves a long-standing spatial disconnect in rDNA biology: the presence of critical, interacting regulatory elements scattered across what was previously considered an unstructured spacer. By defining the start of the unit at the (CTTT)n MMDP-R, the rDNA-GU structurally encapsulates the complete network of cis-regulatory elements required for rRNA transcription into a single, logical upstream domain.

Extensive functional evidence supports the classification of this upstream region as a highly active regulatory hub. Previous studies have identified a spacer promoter located ∼2 kb upstream of the core promoter. Transcripts originating from this site, known as promoter RNA (pRNA), are essential for recruiting the Nucleolar Remodeling Complex (NoRC) to establish the repressive chromatin states that silence rRNA gene transcription (Mayer et al., 2006, 2008; Santoro et al., 2010). Concurrently, tandem enhancer repeats situated between this spacer promoter and the core promoter actively drive transcription through long-range chromatin interactions. (De Winter & Moss, 1986; Tower et al., 1989; Kuhn et al., 1990; Paalman et al., 1995; Caudy & Pikaard, 2002). Although this classic model of repeat-driven enhancement is primarily derived from model organisms like Xenopus and mouse, human rDNA achieves analogous transcriptional enhancement through a distinct topological mechanism. Specifically, a CTCF/cohesin-mediated "enhancer boundary complex" localizes near the human spacer promoter (Herdman et al., 2017; Mars et al., 2018). Rather than relying on repetitive sequences, this complex drives localized 3D chromatin looping, physically bridging the upstream regulatory elements with the core promoter to actively stimulate transcription.

Building upon this topological framework, our microsatellite landscape analysis reveals a critical new dimension to the human rDNA architecture. Strikingly, the most conserved (CTTT)n accumulation peak is not located at the ∼2 kb spacer promoter, but rather at the very beginning of the (CTTT)n MMDP-R, approximately 4 kb upstream of the TSS. This prominent 4 kb peak likely serves as the master structural boundary or "insulator-promoter anchor" of the cis-regulatory domain. We therefore propose a hierarchical control model: the 4 kb (CTTT)n peak acts as a rigid upstream boundary that protects the unit’s regulatory environment from bleeding into the preceding sequence, while the 2 kb region and its localized (CTTT)n tandem clusters act as the proximal "engine" for orchestrating both activation and silencing.

Thus, the (CTTT)n MMDP-R does not merely act as a structural bookend; it serves as the foundational anchor for a comprehensive regulatory cascade. By starting the unit here, our model correctly positions the distal enhancers, the spacer promoter, and the tandem repeats as a contiguous upstream regulatory apparatus that governs the activation and silencing of the adjacent ribosomal genes. The heavy concentration of (CTTT)n repeats in this region likely plays a role in facilitating the unique chromatin architecture or transcription factor recruitment (such as UBF or SL1) required to maintain this complex regulatory environment.

### 4.2. The Downstream (CT)n MDP-CR Functions as a Structural Boundary and Genomic Insulator

While the upstream (CTTT)n MDP-R serves as the regulatory command center for the rDNA-GU, our model designates the downstream (CT)n MDP-CR as its essential structural bookend. It is important to note that this distal boundary is not a monolithic block of simple repeats; rather, it is a massive domain spanning approximately 27,892 bp. Its defining architecture is characterized by three prominent, high-density (CT)n accumulation clusters localized in the middle of the segment, interspersed with less stable microsatellite peaks. Defining the unit’s terminus precisely at this clustered (CT)n landscape clarifies the biophysical mechanics required to maintain stability in a highly transcribed tandem array.

Genomic regions containing concentrated (CT)n repeats—often referred to as polypyrimidine/polypurine (Y/R) tracts—are well documented for their ability to adopt non-B DNA conformations, such as intramolecular triplexes (H-DNA). Foundational biophysical studies have demonstrated that long polypyrimidine tracts readily transition into rigid triplex DNA structures when subjected to high negative supercoiling, even at a neutral pH and physiological ionic strength in the presence of Mg²□ (Sakamoto et al., 1996).

Beyond triplex formation, these downstream sequence landscapes are heavily implicated in the formation of scheduled, regulatory R-loops (complexes comprising a DNA:RNA hybrid and displaced single-stranded DNA) (Smirnov et al., 2021). Recent single-molecule footprinting has revealed that R-loops are non-randomly distributed across human rDNA and play a critical role in nucleolar organization. Strikingly, recent data indicate that antisense transcription by RNA polymerase II generates a protective R-loop shield within the intergenic regions. This shield physically protects nucleolar organization by suppressing disruptive, unscheduled non-coding RNAs transcribed by RNA polymerase I.

Therefore, the presence of these three massive (CT)n clusters at the distal boundary of the rDNA-GU is highly functional. In the context of the nucleolus, the (CT)n MDP-CR likely operates as an inducible, multi-layered biophysical barrier. By facilitating the formation of both supercoiling-induced triplexes and regulatory "R-loop shields," these clusters provide the exact structural roadblocks needed to stall replication forks, prevent transcriptional read-through from the preceding core, and physically insulate adjacent rDNA units. By resolving the former intergenic spacer into distinct regulatory (upstream) and structural (downstream) compartments, the rDNA-GU model accurately reflects the biophysical realities of nucleolar organization.

### 4.3. Distinct Biophysical Signatures of Microsatellites Drive 3D Regulatory Architecture

The distinct spatial distributions of (CT)n and (CTTT)n tracts within the human rDNA intergenic spacer strongly suggest that their regulatory roles are fundamentally rooted in their divergent biophysical properties. While both are pyrimidine-rich, it is crucial to distinguish the thermodynamic and mechanical differences driven by their varying GC contents. Standard dinucleotide (CT)n arrays maintain a 50% GC content and are well-characterized for their structural rigidity and propensity to adopt non-B DNA conformations, such as H-DNA (intramolecular triplexes), under physiological supercoiling. In stark contrast, the tetranucleotide (CTTT)n motif possesses a significantly lower GC content of 25%. This substantial reduction in G-C base pairing fundamentally alters the double helix’s mechanical properties. The T-rich nature of the (CTTT)n tract lowers the local melting temperature, thereby increasing DNA "breathability" (the propensity for localized unwinding) and imparting a distinct intrinsic helical flexibility and curvature.

We hypothesize that these precise topological features, dictated by the specific GC-to-AT ratio, act as a structural fingerprint for the master boundary. Rather than functioning merely as sequence-specific binding sites, the localized DNA curvature and breathability provided by the prominent 4 kb (CTTT)n peak likely furnish the exact physical topology required to exclude nucleosomes or efficiently anchor architectural protein complexes like CTCF and Cohesin. Ultimately, the utilization of specific (CTTT)n arrays over generic (CT)n repeats highlights a sophisticated layer of genomic design: a shift from simple sequence variation to targeted structural modulation, which precisely dictates the 3D spatial organization and functional partitioning of the human rDNA regulatory hub.

## Abbreviations

STRs: Short Tandem Repeats
HMDPs: High Microsatellites Density Peaks
MMDPs: Middle Microsatellites Density Peaks
LMDPs: Low Microsatellites Density Peaks
MDPs: Microsatellite Density Peaks
DSU: Duplication Segment Unit
5′-ETS: 5′ External Transcribed Spacer
ITS1: Internal Transcribed Spacer 1
IGS: Intergenic Spacer
rDNA-GUs: rDNA Gene Units
NORs: Nucleolar Organizer Regions
(CTTT)n MDPs-R: (CTTT)n MDPs Region
(CT)n MDPs-CR: (CT)n MDPs Cluster Region
Co-0(CT/CTTT)n MDP-Rs: Common Zero (CT/CTTT)n Microsatellite Density Peak Regions
Co-0(CT/CTTT)n MDP-BRs: Common Zero (CT/CTTT)n Microsatellite Density Peak Big Regions

## Declarations

### Ethics approval and consent to participate

Not applicable.

### Consent for publication

Not applicable.

### Availability of data and materials

The microsatellite density landscapes of 24 chromosomes of T2T-CHM13, including specific tracks for (CT)n/(CTTT)n patterns, can be accessed at http://genome.ucsc.edu/s/zhongyangtan/CHM13v2.1.

All data generated or analysed during this study are included in this published article and its supplementary information files.

### Authors’ contributions

Z.T. designed and directed this study. J.S and S.T. prepared the manuscript. J.S., S.T., Y.X, J.Q and H.X. developed the statistical method and programs. J.S., S.T., Y.X, J.Q and H.X. performed the data analysis, Z.T. edited this manuscript. All authors read and approved the final version of the manuscript.

