## Supplementary figures and images for "Clarified an rDNA Gene Unit Pattern with (CTTT)n and (CT)n Microsatellites Aggregation Ahead of and Behind the Gene in Human Genome"

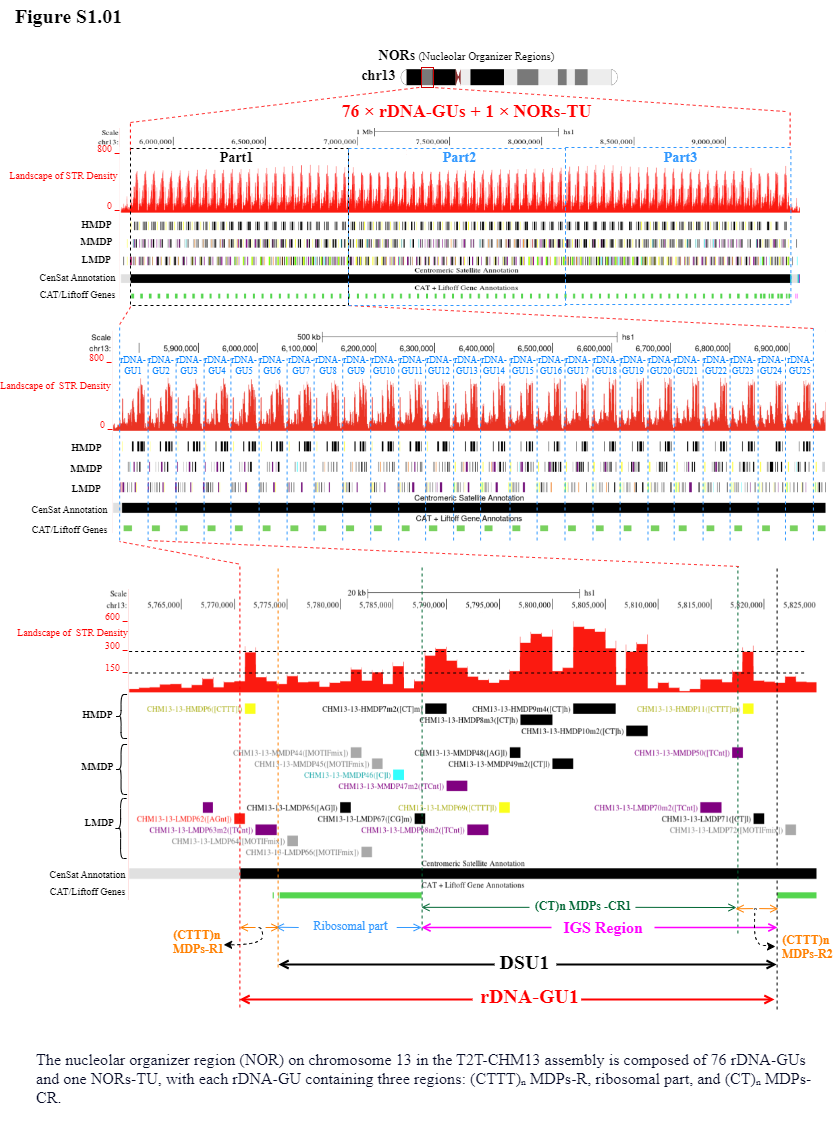

Supplement: Supplementary Table and Figure [file 713381_file03.zip › Fig and Table of Manuscript_20260322/Supplementary Figure/Supplementary Figure1_NORs in the T2T-CHM13 assembly are composed of multiple rDNA-GUs and a single NORs-TU/S1.01.png]

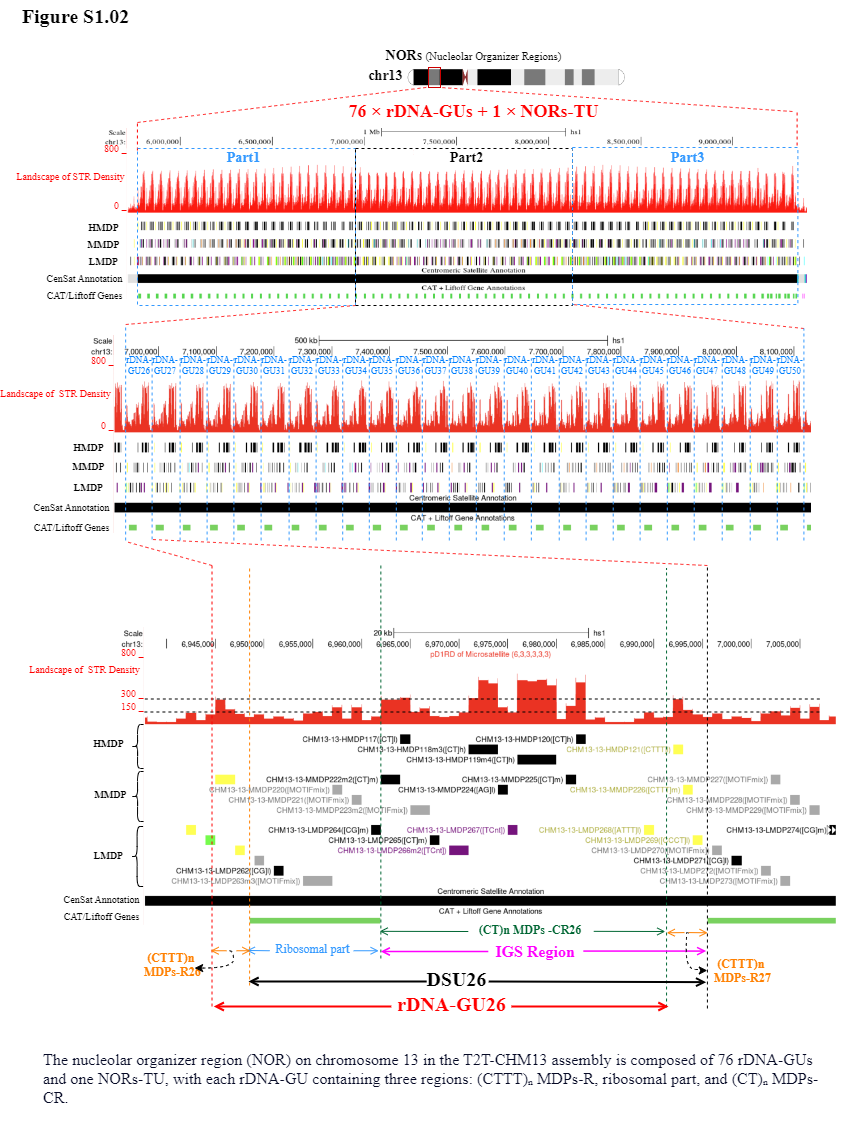

Supplement: Supplementary Table and Figure [file 713381_file03.zip › Fig and Table of Manuscript_20260322/Supplementary Figure/Supplementary Figure1_NORs in the T2T-CHM13 assembly are composed of multiple rDNA-GUs and a single NORs-TU/S1.02..png]

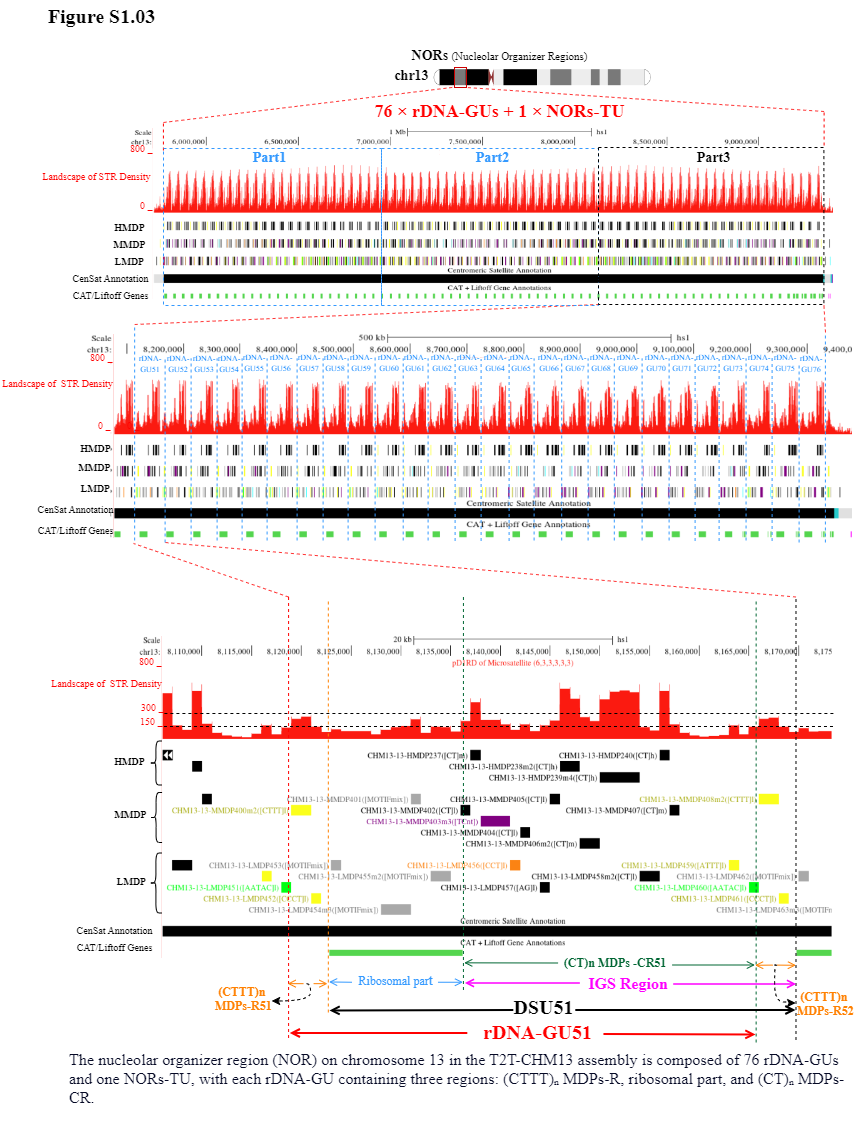

Supplement: Supplementary Table and Figure [file 713381_file03.zip › Fig and Table of Manuscript_20260322/Supplementary Figure/Supplementary Figure1_NORs in the T2T-CHM13 assembly are composed of multiple rDNA-GUs and a single NORs-TU/S1.03.png]

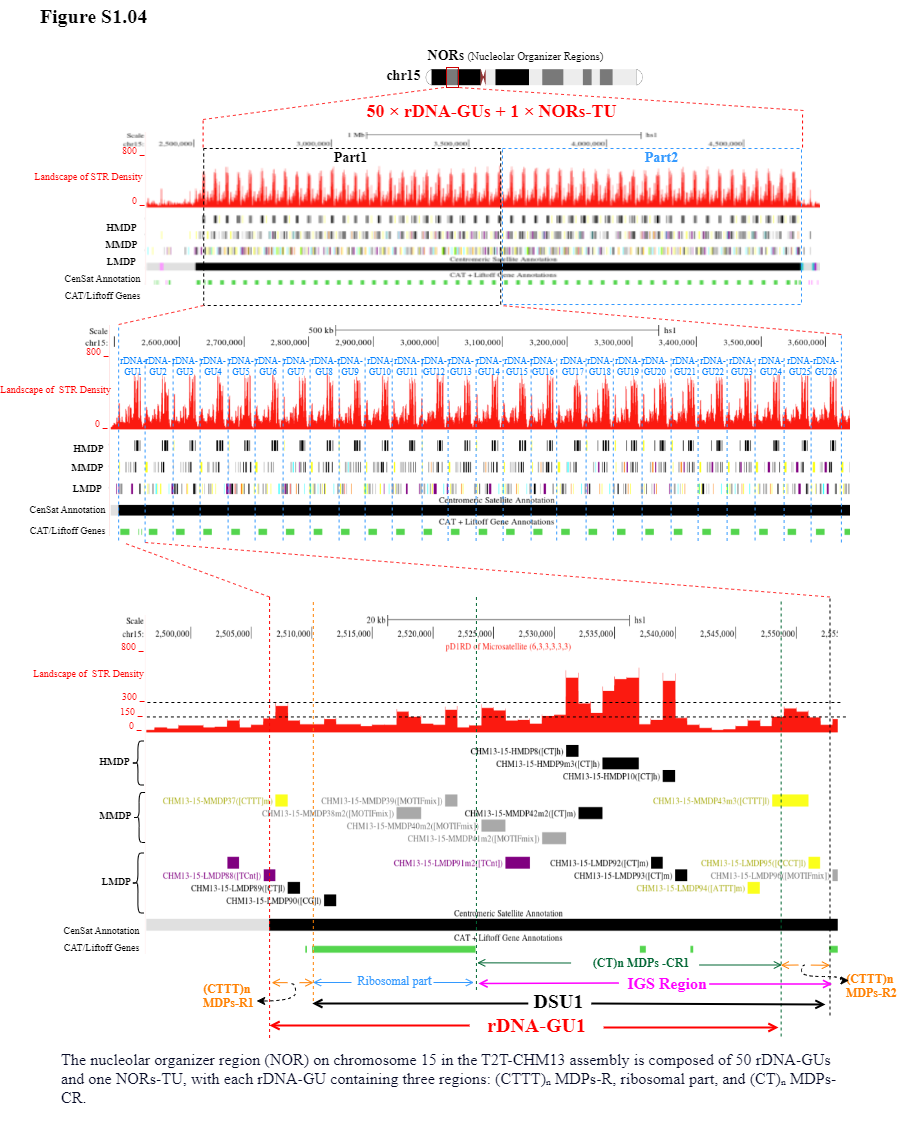

Supplement: Supplementary Table and Figure [file 713381_file03.zip › Fig and Table of Manuscript_20260322/Supplementary Figure/Supplementary Figure1_NORs in the T2T-CHM13 assembly are composed of multiple rDNA-GUs and a single NORs-TU/S1.04.png]

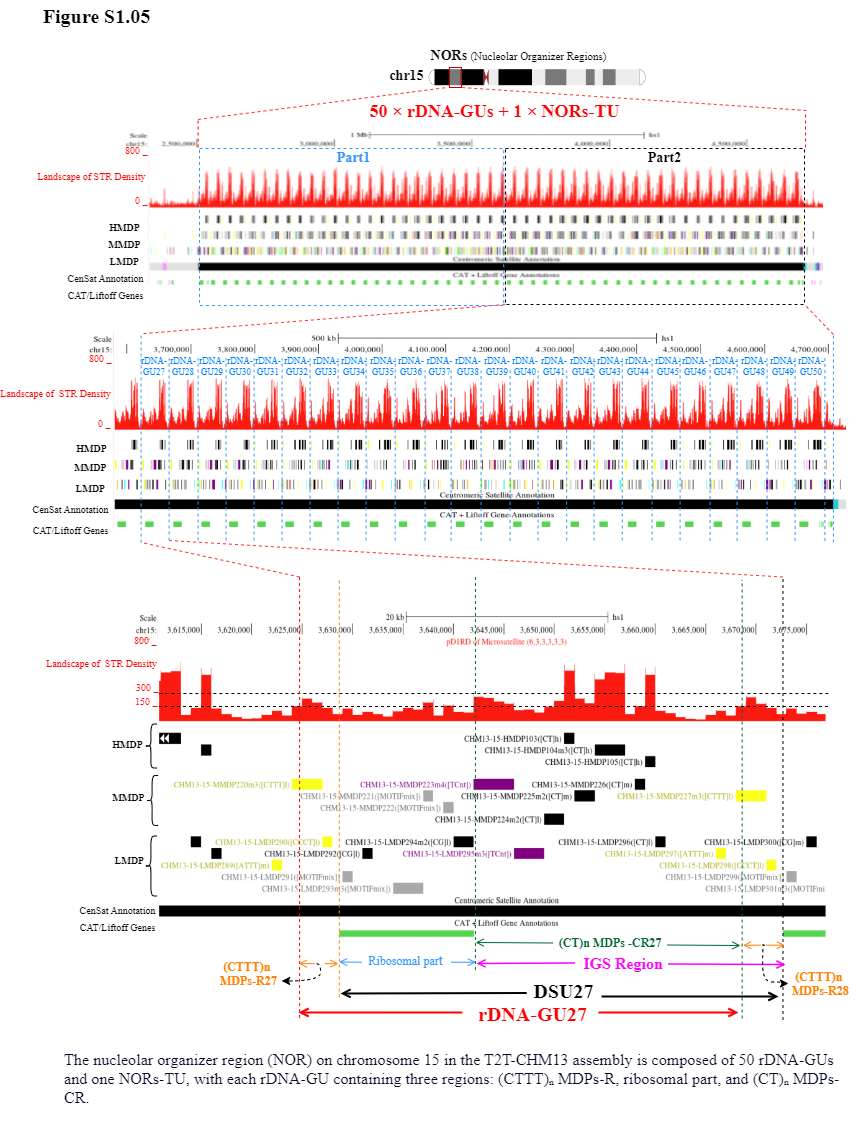

Supplement: Supplementary Table and Figure [file 713381_file03.zip › Fig and Table of Manuscript_20260322/Supplementary Figure/Supplementary Figure1_NORs in the T2T-CHM13 assembly are composed of multiple rDNA-GUs and a single NORs-TU/S1.05.png]

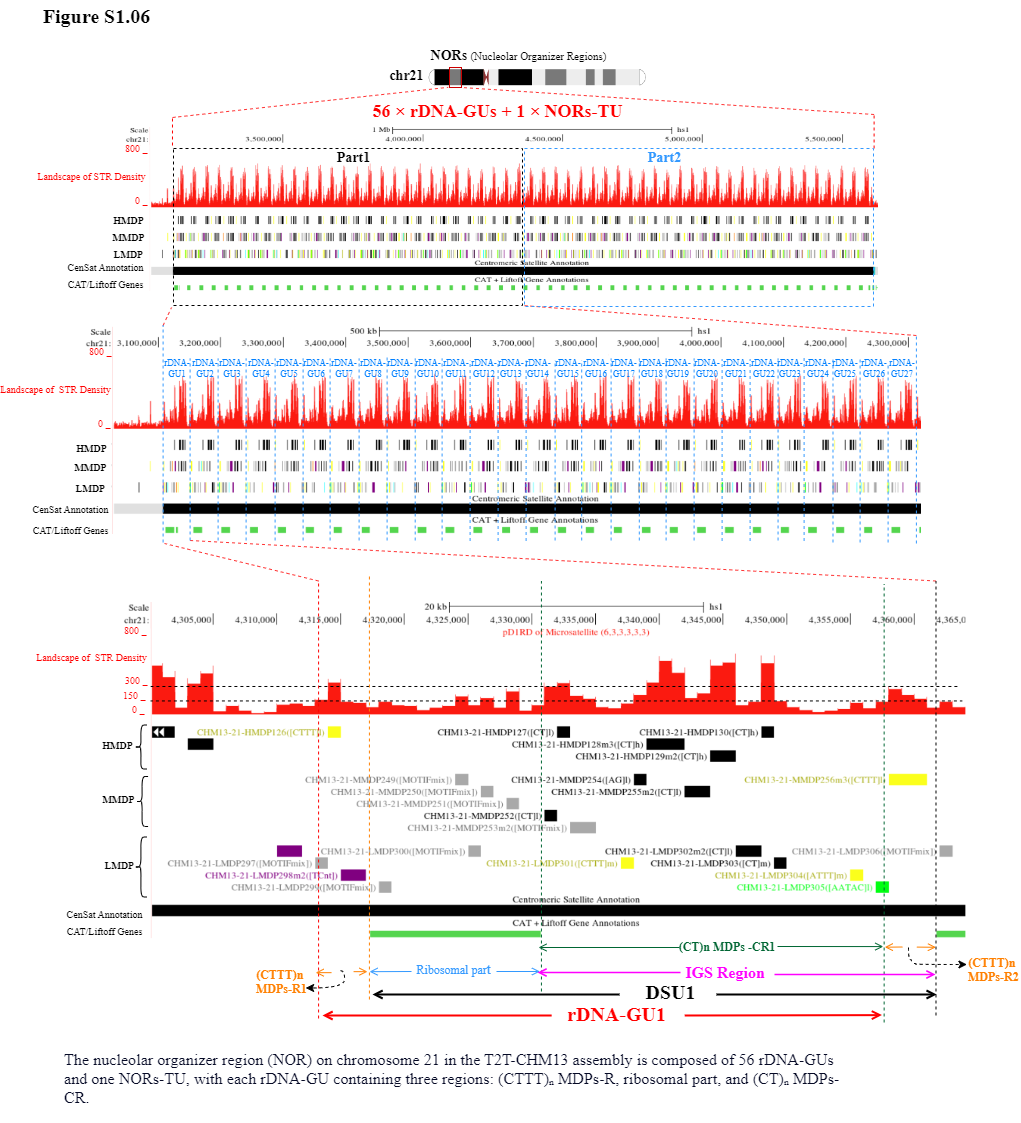

Supplement: Supplementary Table and Figure [file 713381_file03.zip › Fig and Table of Manuscript_20260322/Supplementary Figure/Supplementary Figure1_NORs in the T2T-CHM13 assembly are composed of multiple rDNA-GUs and a single NORs-TU/S1.06.png]

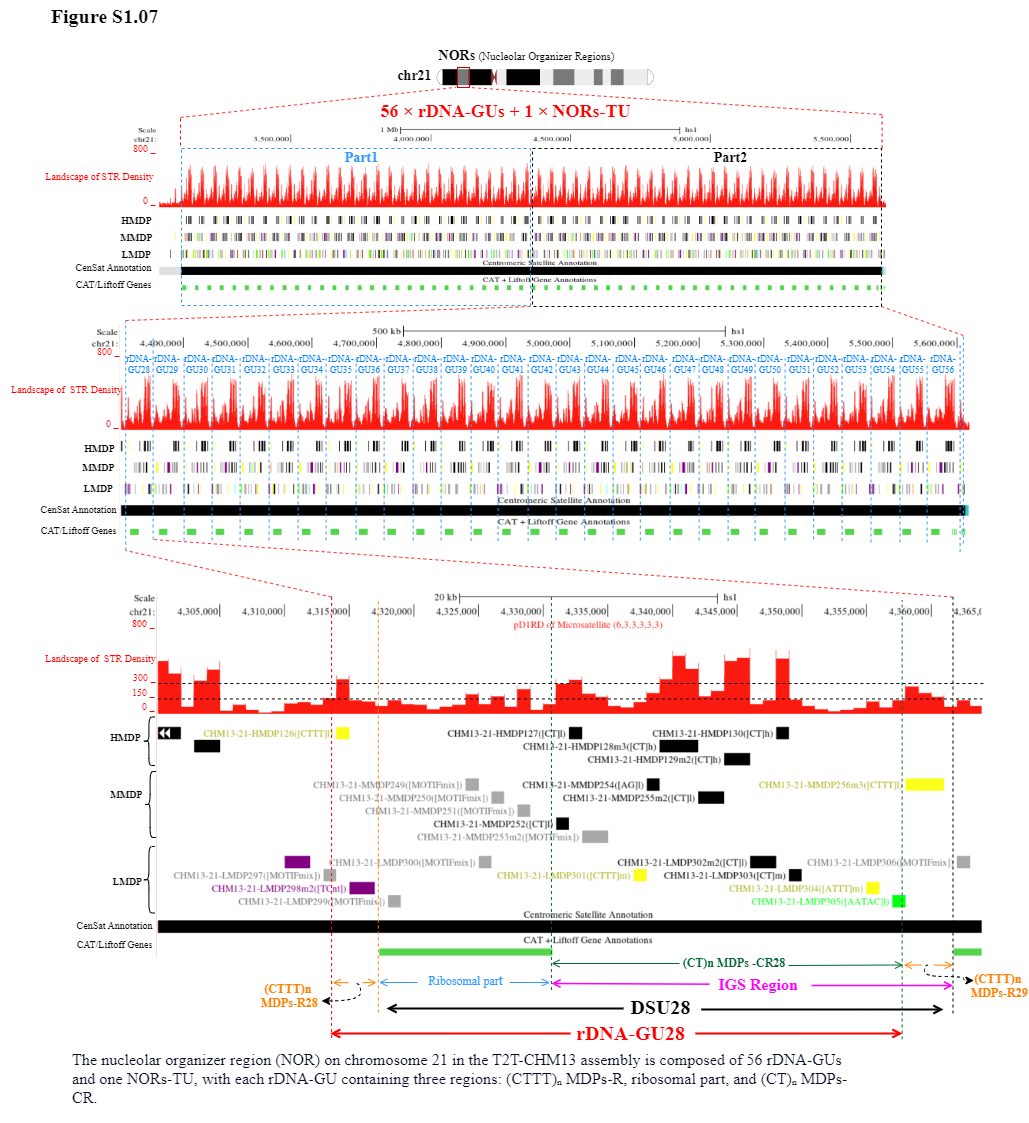

Supplement: Supplementary Table and Figure [file 713381_file03.zip › Fig and Table of Manuscript_20260322/Supplementary Figure/Supplementary Figure1_NORs in the T2T-CHM13 assembly are composed of multiple rDNA-GUs and a single NORs-TU/S1.07.png]

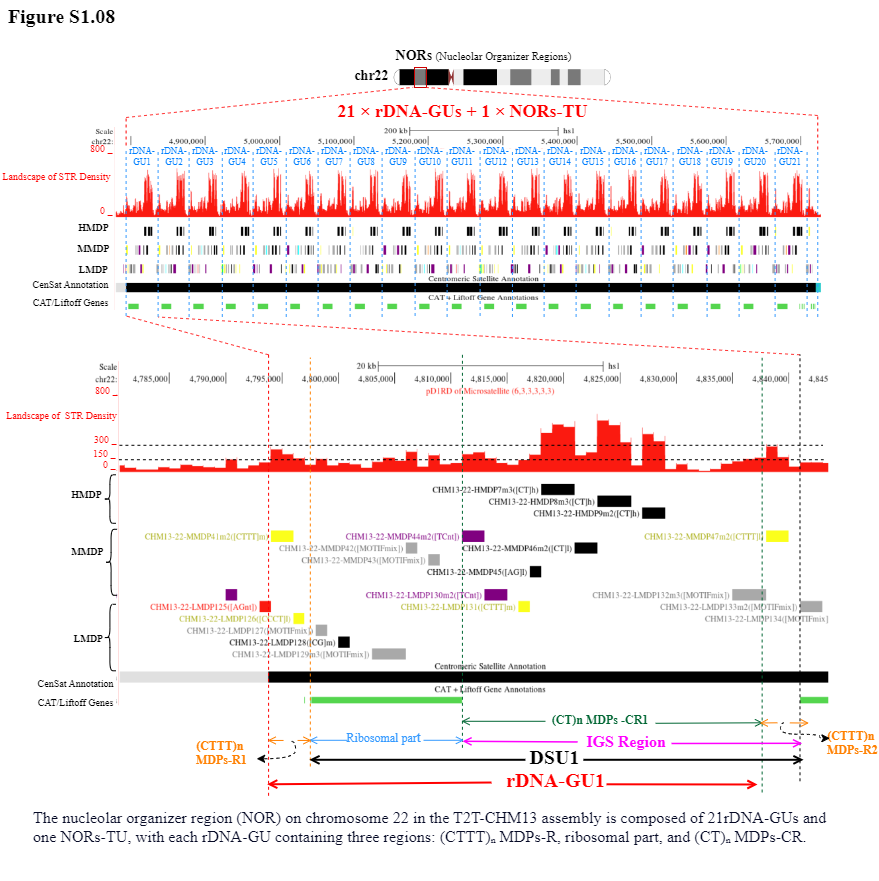

Supplement: Supplementary Table and Figure [file 713381_file03.zip › Fig and Table of Manuscript_20260322/Supplementary Figure/Supplementary Figure1_NORs in the T2T-CHM13 assembly are composed of multiple rDNA-GUs and a single NORs-TU/S1.08.png]

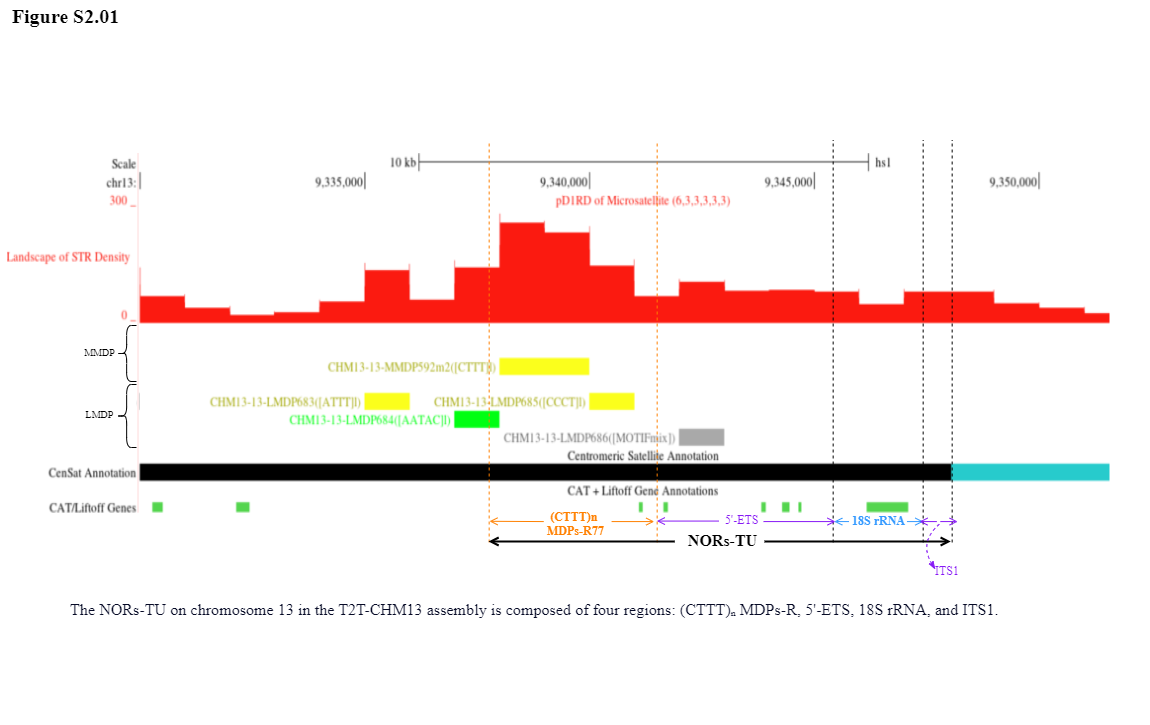

Supplement: Supplementary Table and Figure [file 713381_file03.zip › Fig and Table of Manuscript_20260322/Supplementary Figure/Supplementary Figure2_NORs-TUs on human acrocentric chromosomes 13, 14, 15, 21, and 22 in the T2T-CHM13 assembly/S2.01.png]

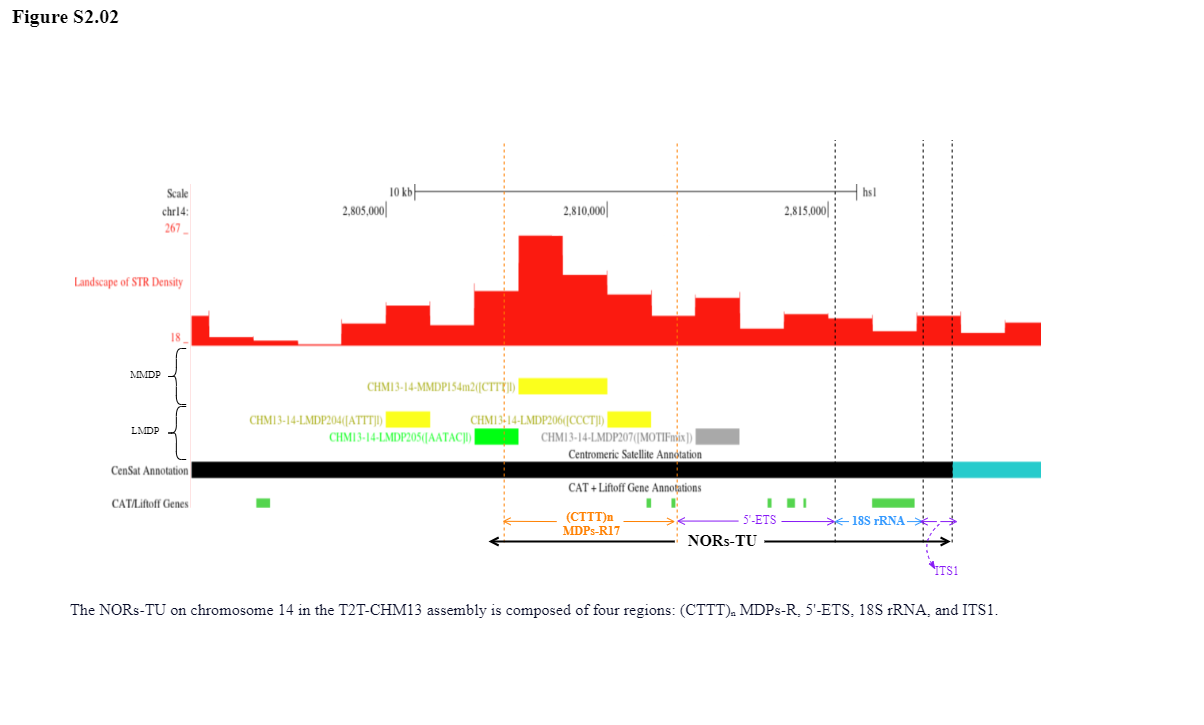

Supplement: Supplementary Table and Figure [file 713381_file03.zip › Fig and Table of Manuscript_20260322/Supplementary Figure/Supplementary Figure2_NORs-TUs on human acrocentric chromosomes 13, 14, 15, 21, and 22 in the T2T-CHM13 assembly/S2.02.png]

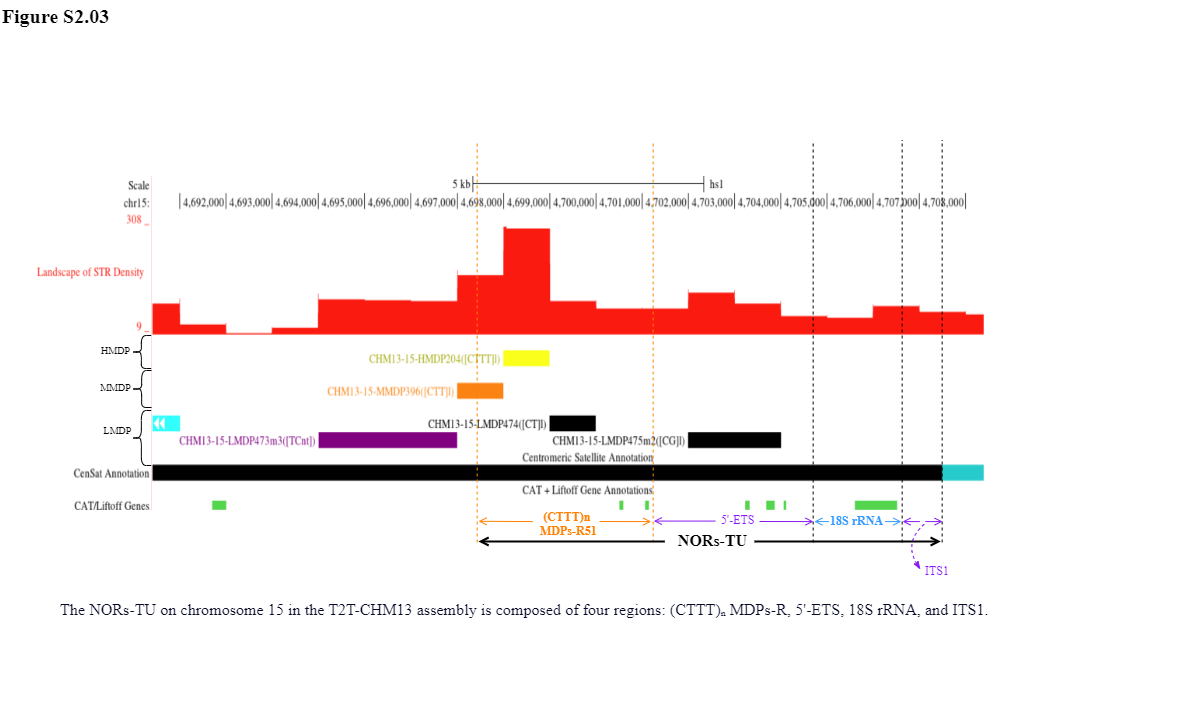

Supplement: Supplementary Table and Figure [file 713381_file03.zip › Fig and Table of Manuscript_20260322/Supplementary Figure/Supplementary Figure2_NORs-TUs on human acrocentric chromosomes 13, 14, 15, 21, and 22 in the T2T-CHM13 assembly/S2.03.png]

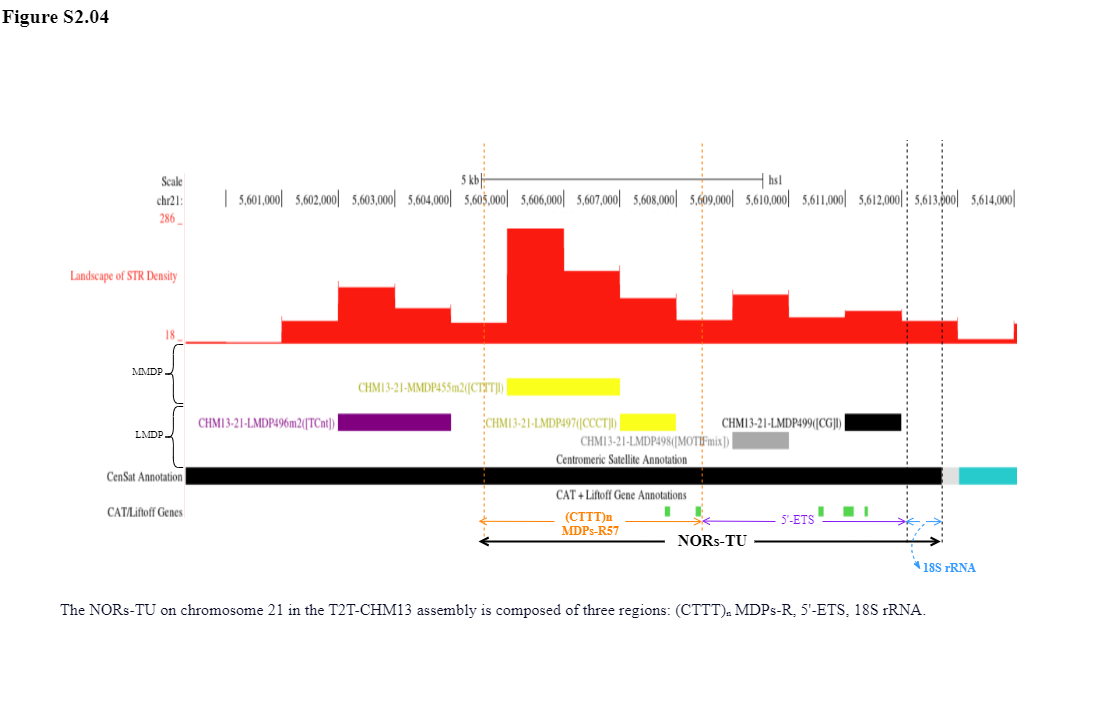

Supplement: Supplementary Table and Figure [file 713381_file03.zip › Fig and Table of Manuscript_20260322/Supplementary Figure/Supplementary Figure2_NORs-TUs on human acrocentric chromosomes 13, 14, 15, 21, and 22 in the T2T-CHM13 assembly/S2.04.png]

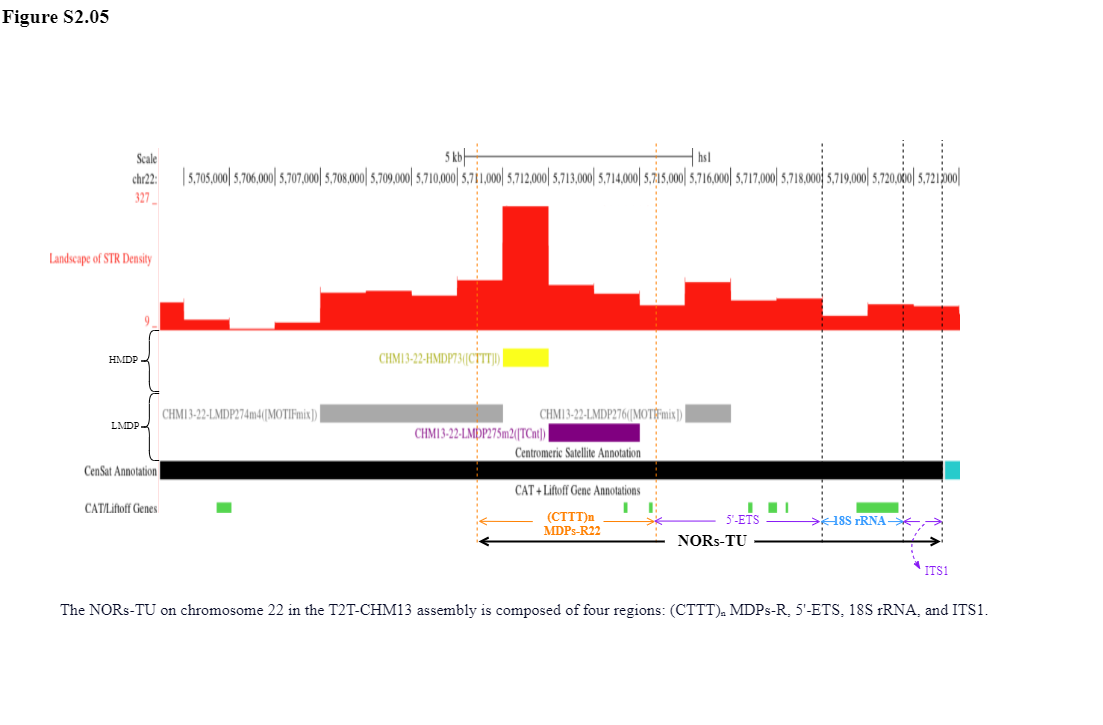

Supplement: Supplementary Table and Figure [file 713381_file03.zip › Fig and Table of Manuscript_20260322/Supplementary Figure/Supplementary Figure2_NORs-TUs on human acrocentric chromosomes 13, 14, 15, 21, and 22 in the T2T-CHM13 assembly/S2.05.png]

### Fig S3.Identity Comparison of 214 (CTTT)n MDPs-R

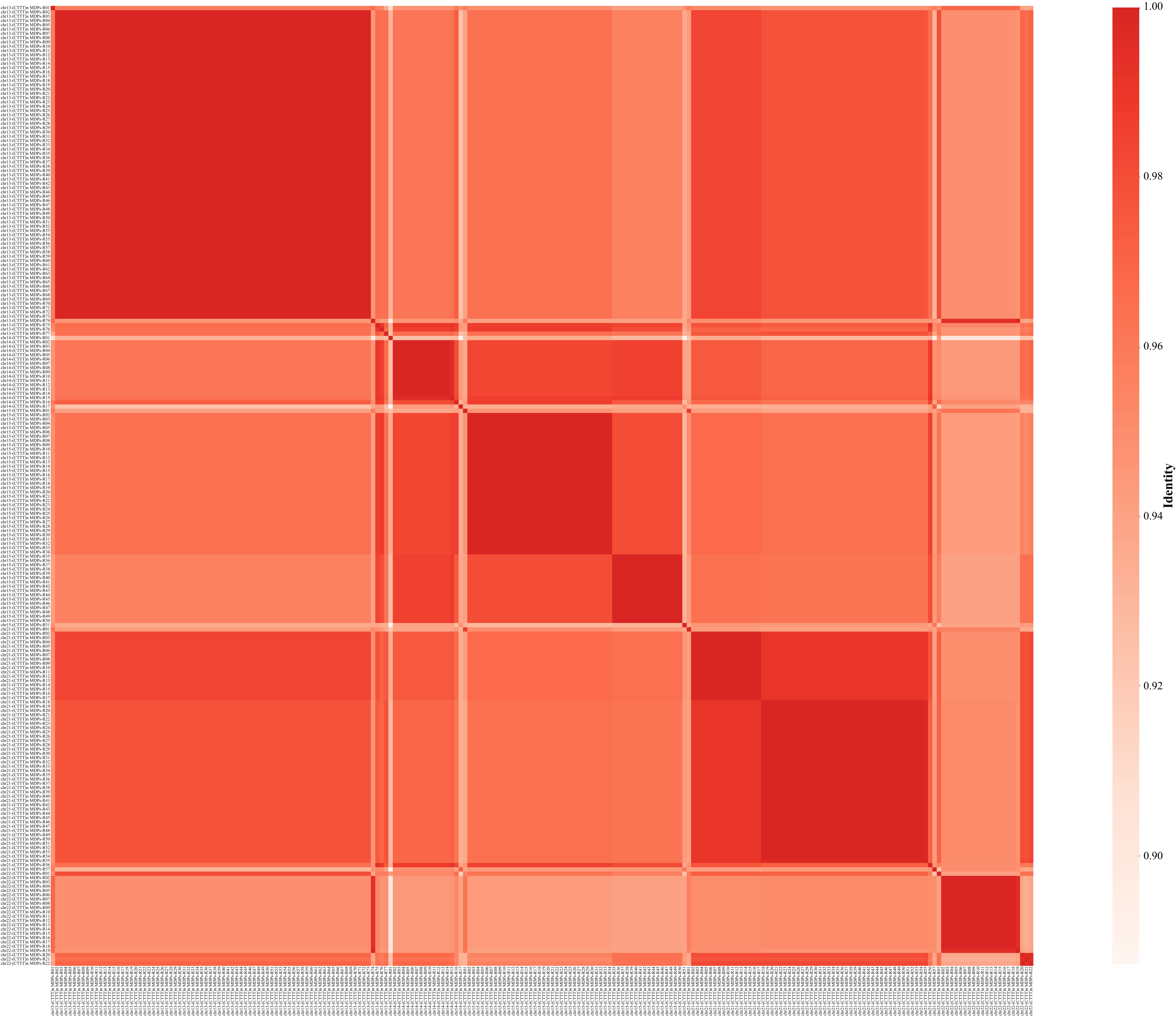

Supplement: Supplementary Table and Figure [file 713381_file03.zip › Fig and Table of Manuscript_20260322/Supplementary Figure/Supplementary Figure3.Identity Comparison of 224 (CTTT)n MDPs-R.pdf]

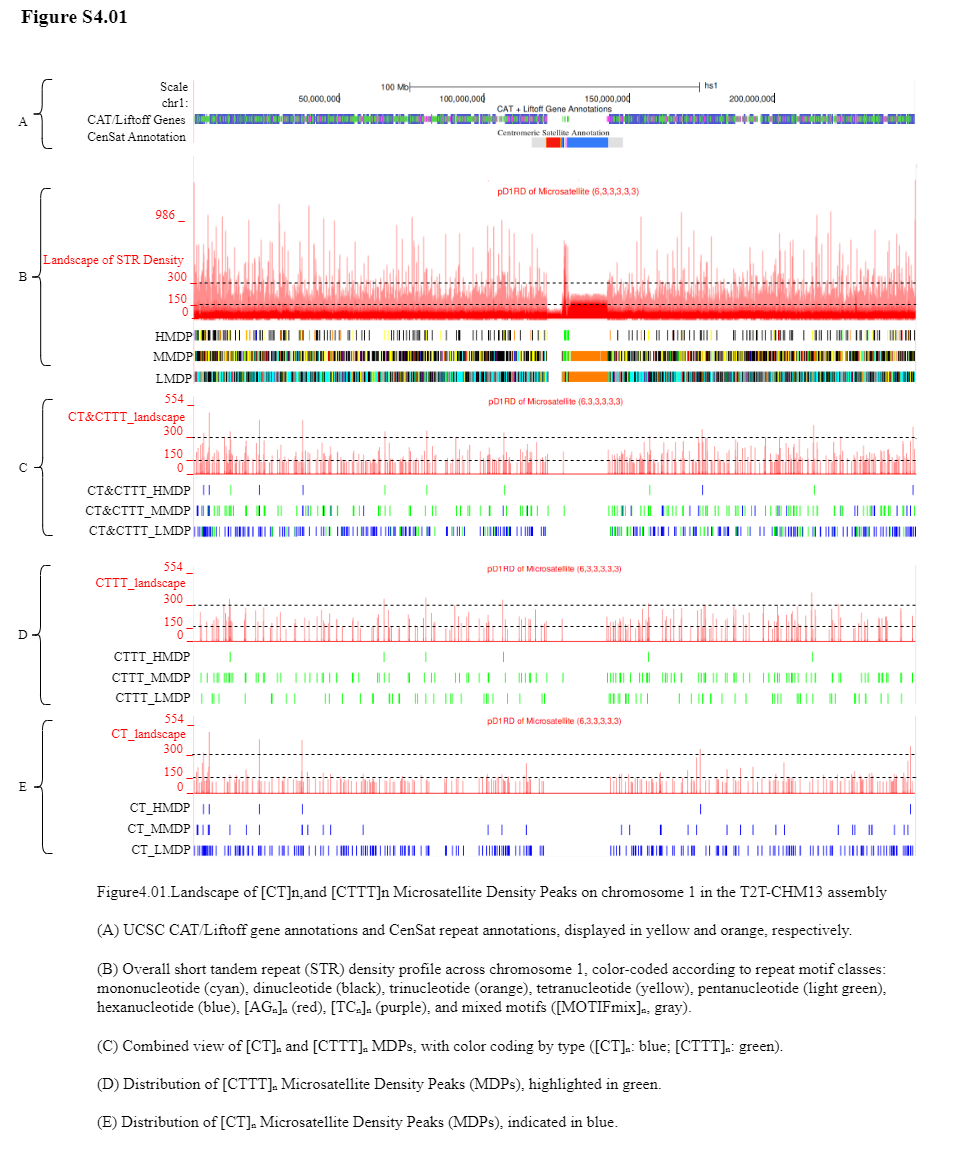

Supplement: Supplementary Table and Figure [file 713381_file03.zip › Fig and Table of Manuscript_20260322/Supplementary Figure/Supplementary Figure4_Landscape of [CT]n,and [CTTT]n Microsatellite Density Peaks in the T2T-CHM13 assembly/S4.01.png]

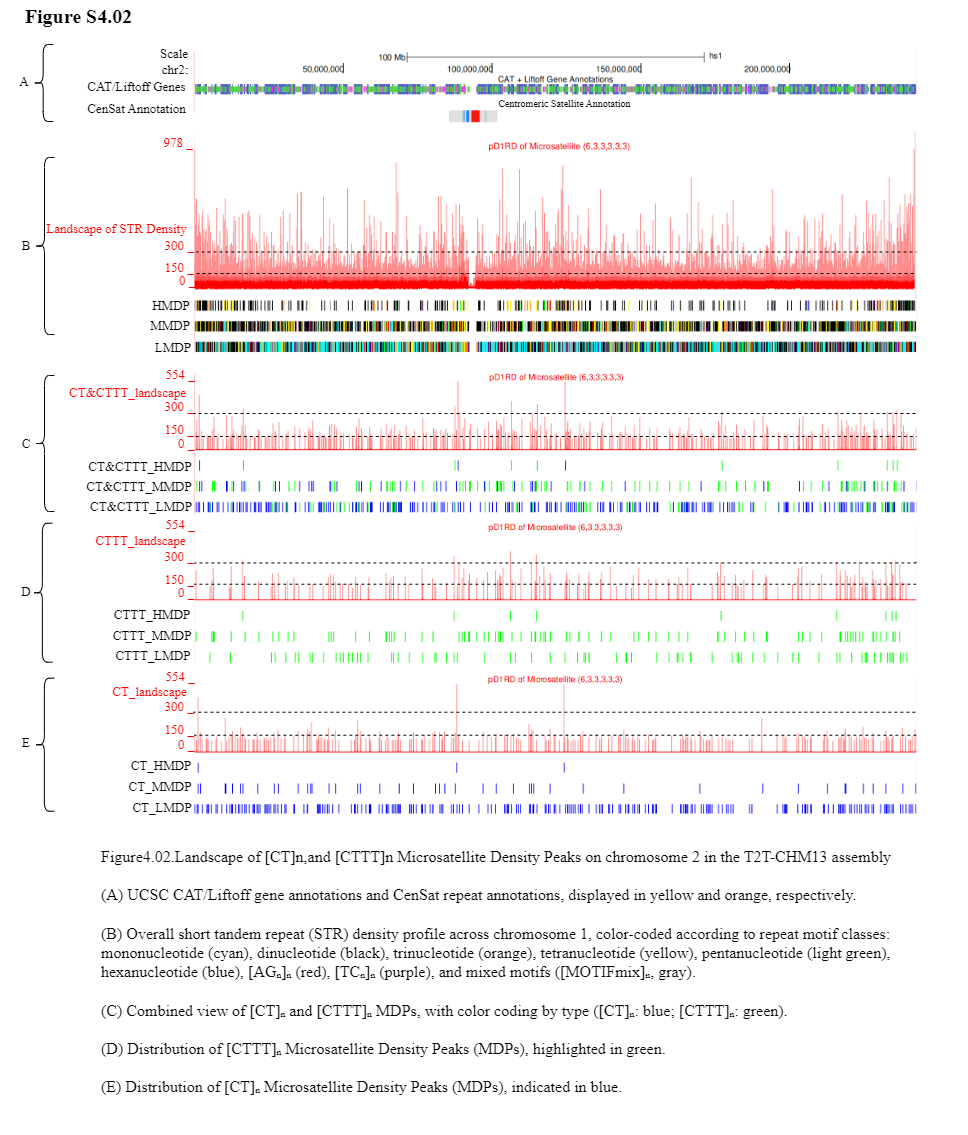

Supplement: Supplementary Table and Figure [file 713381_file03.zip › Fig and Table of Manuscript_20260322/Supplementary Figure/Supplementary Figure4_Landscape of [CT]n,and [CTTT]n Microsatellite Density Peaks in the T2T-CHM13 assembly/S4.02.png]

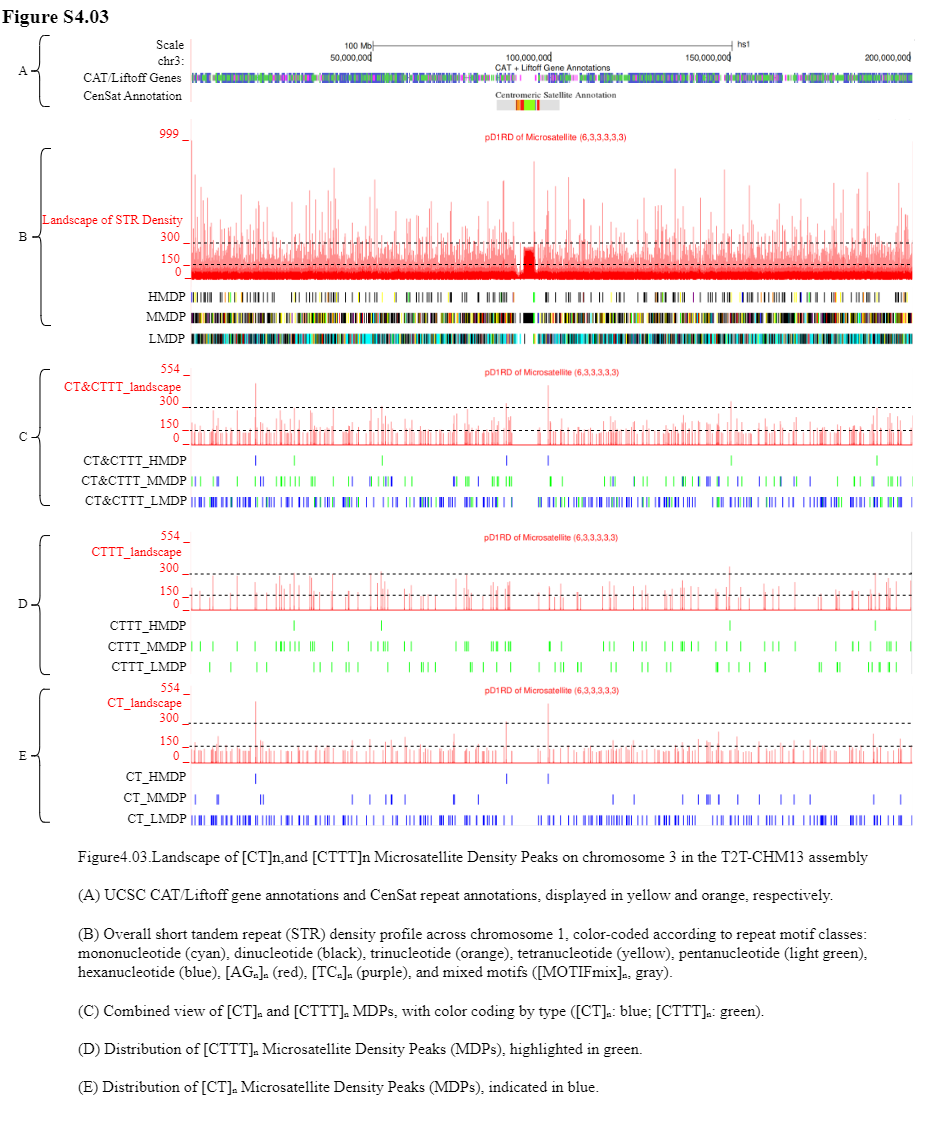

Supplement: Supplementary Table and Figure [file 713381_file03.zip › Fig and Table of Manuscript_20260322/Supplementary Figure/Supplementary Figure4_Landscape of [CT]n,and [CTTT]n Microsatellite Density Peaks in the T2T-CHM13 assembly/S4.03.png]

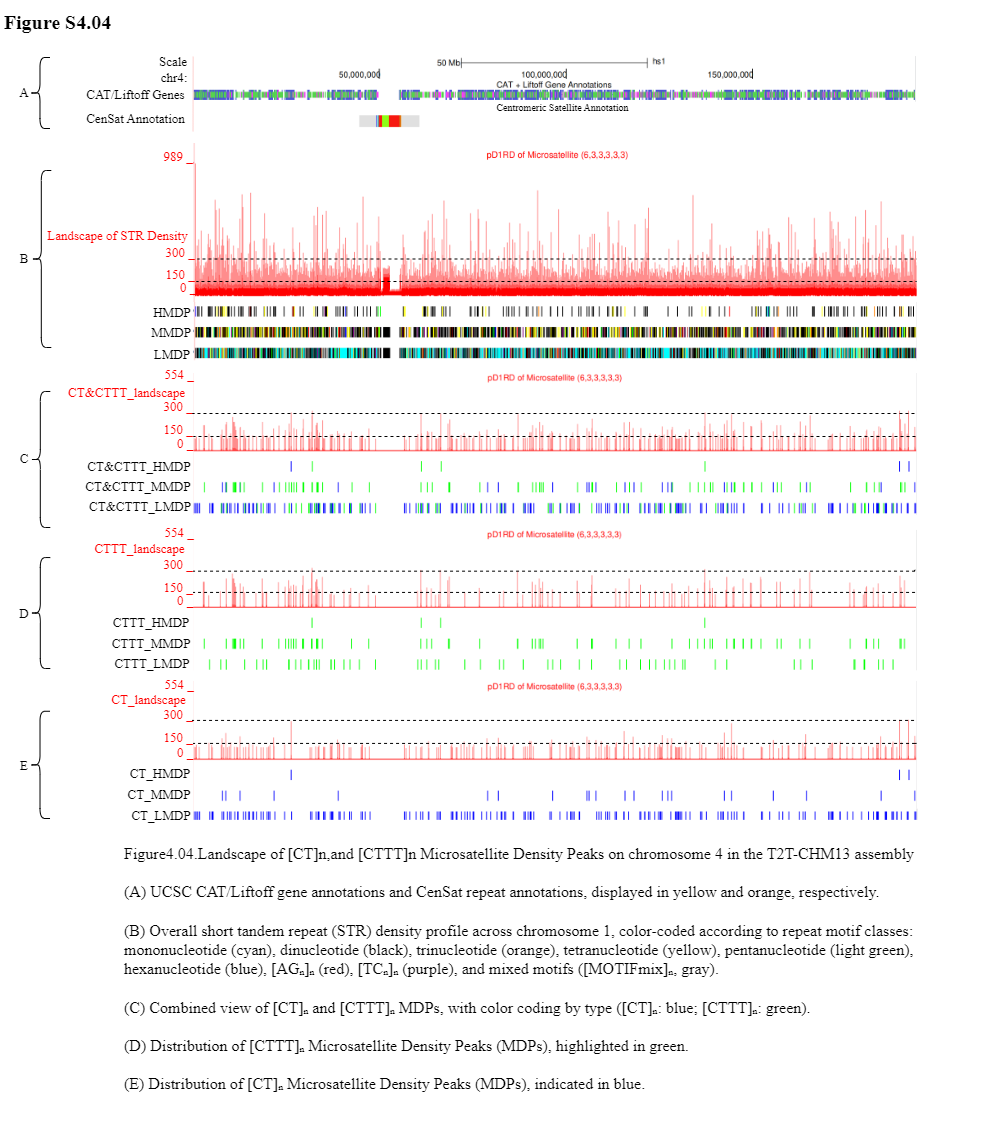

Supplement: Supplementary Table and Figure [file 713381_file03.zip › Fig and Table of Manuscript_20260322/Supplementary Figure/Supplementary Figure4_Landscape of [CT]n,and [CTTT]n Microsatellite Density Peaks in the T2T-CHM13 assembly/S4.04.png]

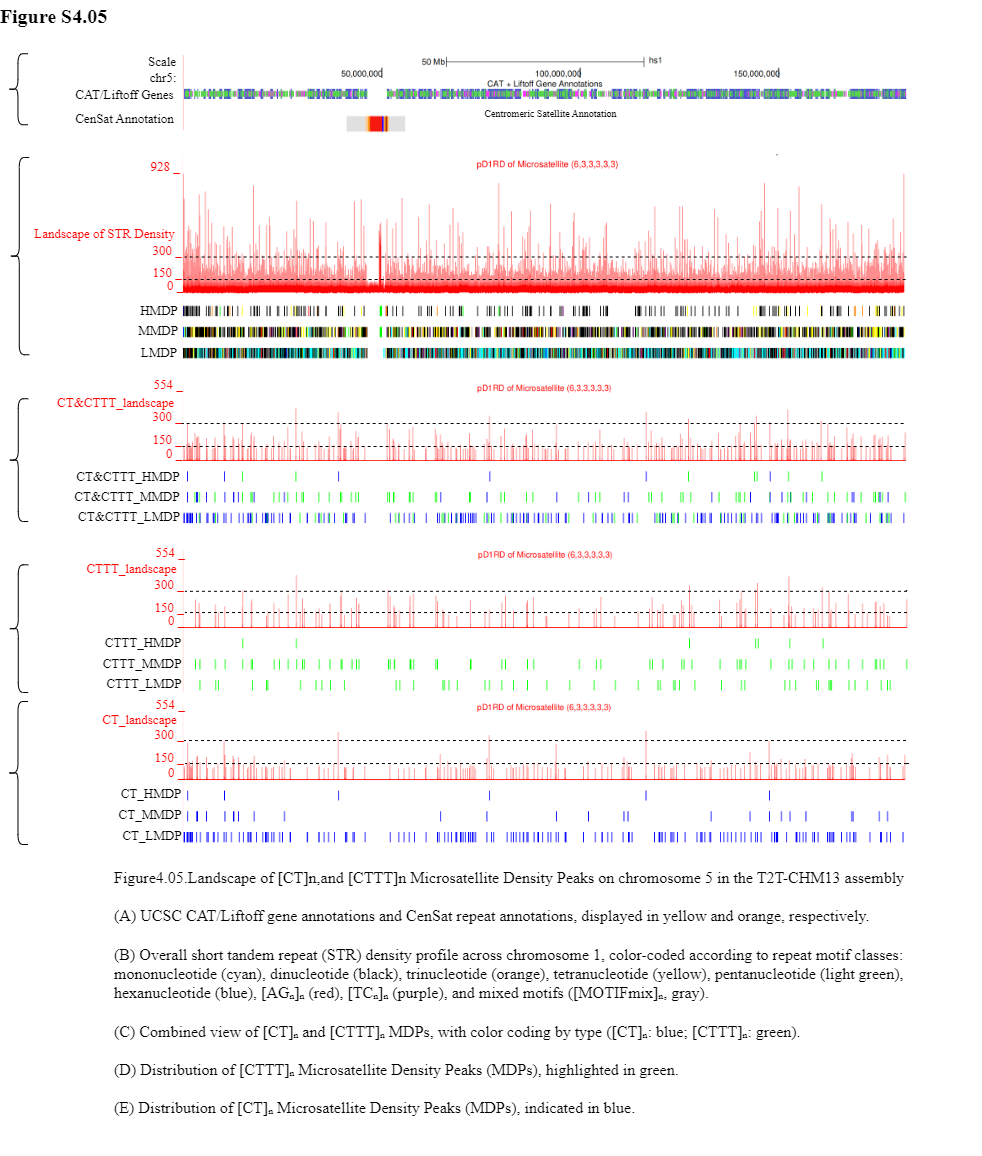

Supplement: Supplementary Table and Figure [file 713381_file03.zip › Fig and Table of Manuscript_20260322/Supplementary Figure/Supplementary Figure4_Landscape of [CT]n,and [CTTT]n Microsatellite Density Peaks in the T2T-CHM13 assembly/S4.05.png]

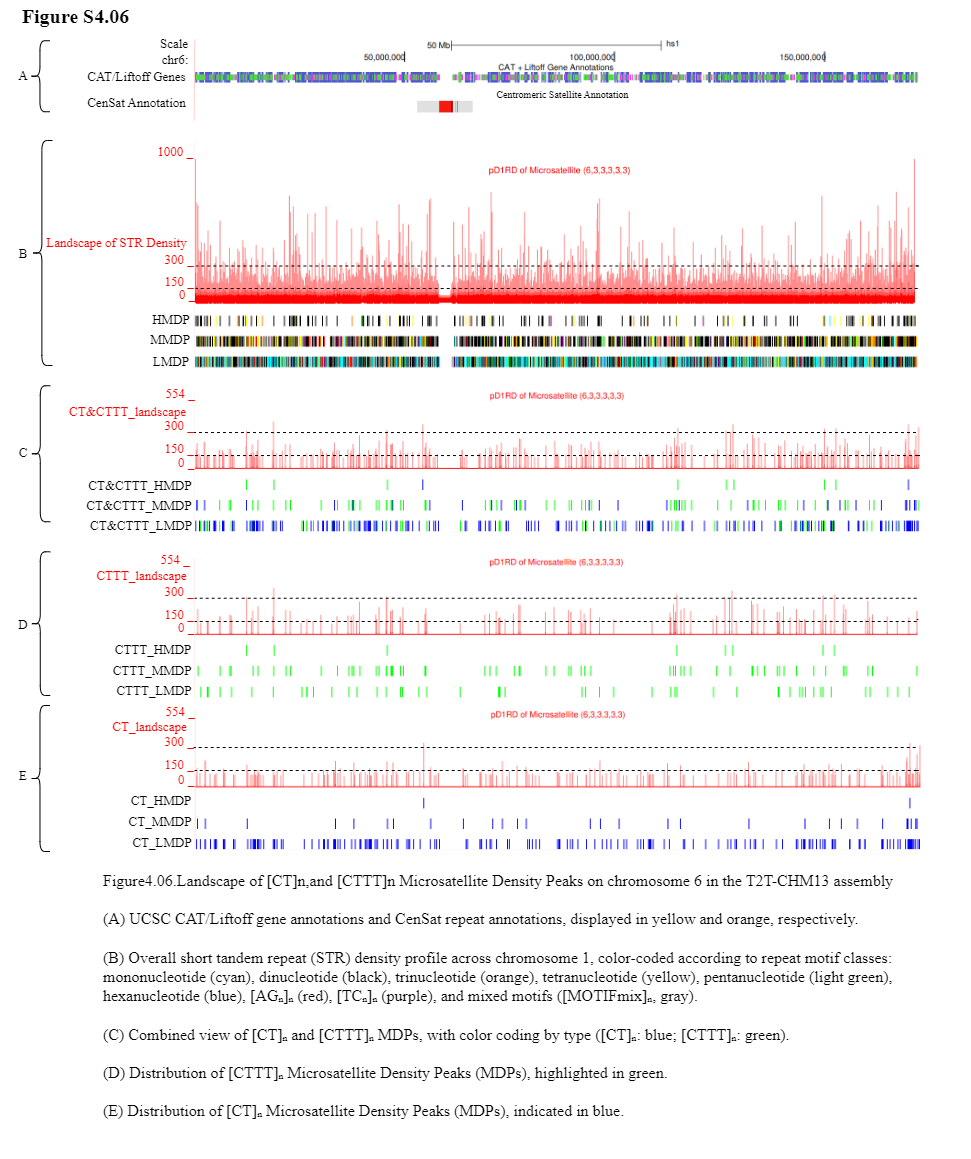

Supplement: Supplementary Table and Figure [file 713381_file03.zip › Fig and Table of Manuscript_20260322/Supplementary Figure/Supplementary Figure4_Landscape of [CT]n,and [CTTT]n Microsatellite Density Peaks in the T2T-CHM13 assembly/S4.06.png]

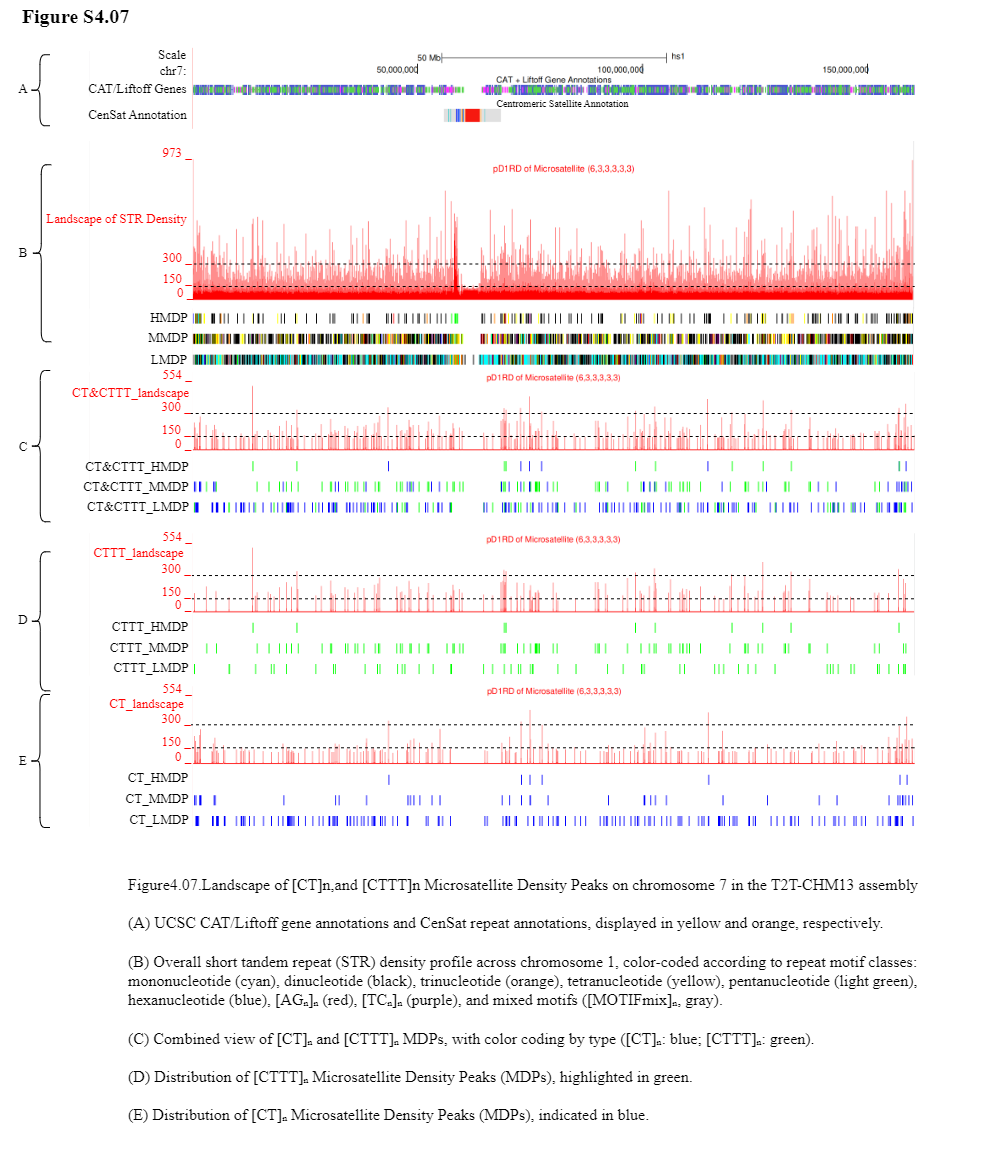

Supplement: Supplementary Table and Figure [file 713381_file03.zip › Fig and Table of Manuscript_20260322/Supplementary Figure/Supplementary Figure4_Landscape of [CT]n,and [CTTT]n Microsatellite Density Peaks in the T2T-CHM13 assembly/S4.07.png]

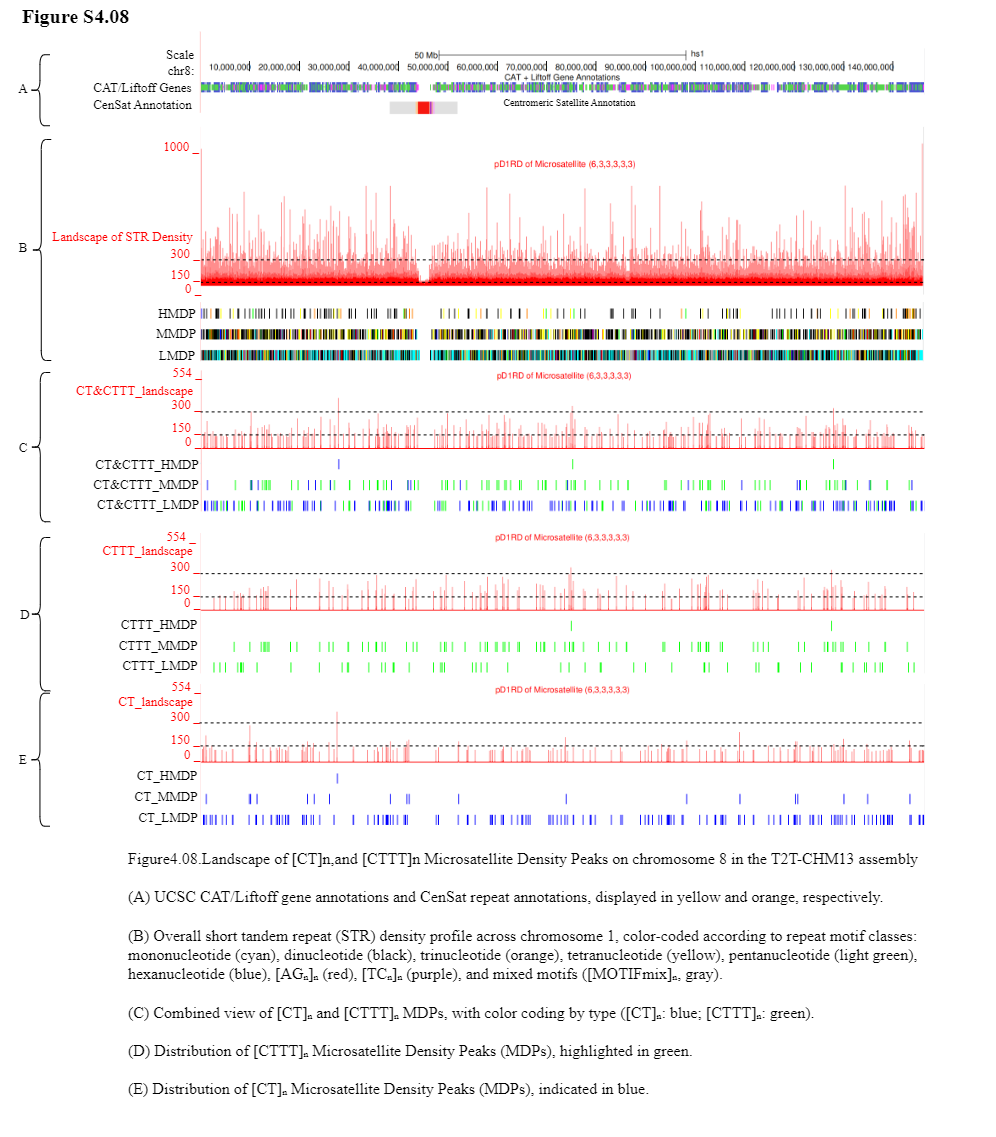

Supplement: Supplementary Table and Figure [file 713381_file03.zip › Fig and Table of Manuscript_20260322/Supplementary Figure/Supplementary Figure4_Landscape of [CT]n,and [CTTT]n Microsatellite Density Peaks in the T2T-CHM13 assembly/S4.08.png]

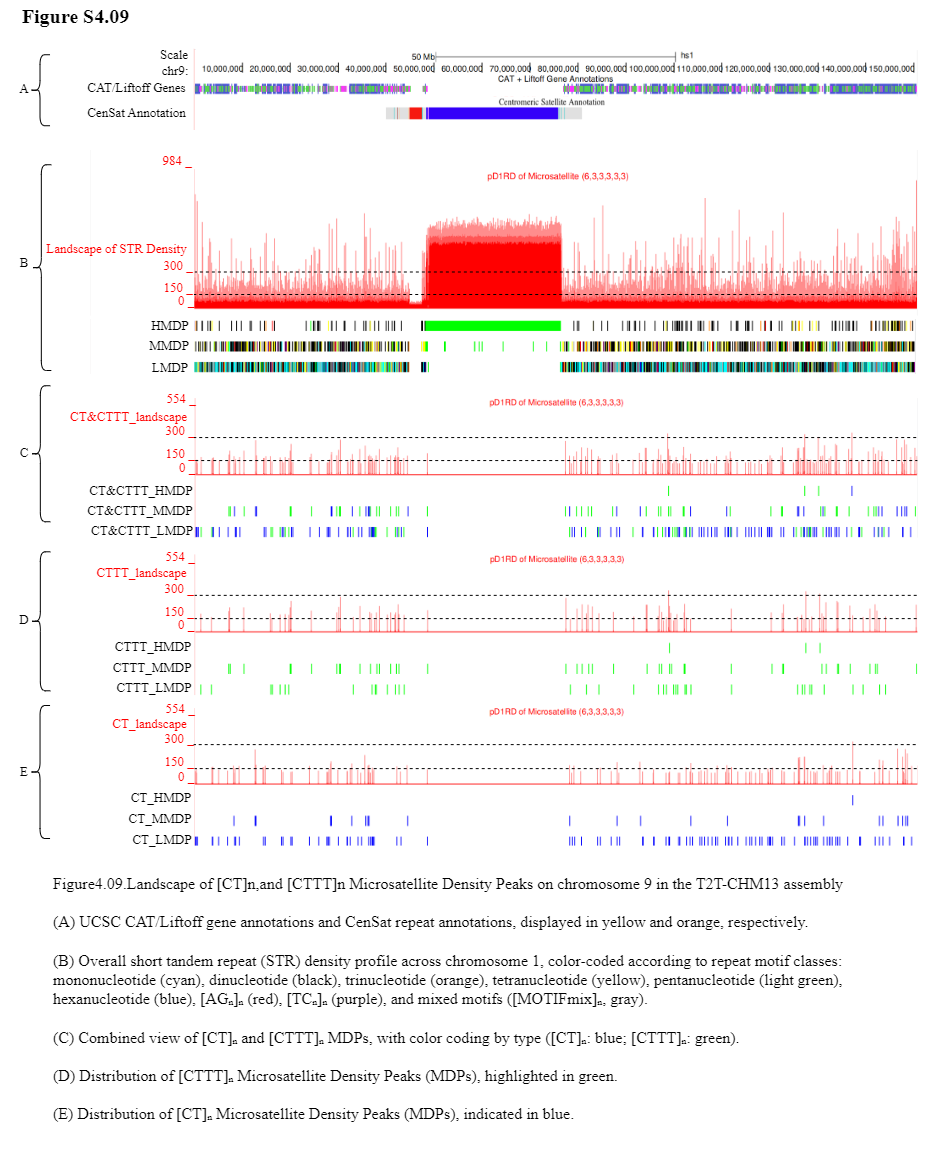

Supplement: Supplementary Table and Figure [file 713381_file03.zip › Fig and Table of Manuscript_20260322/Supplementary Figure/Supplementary Figure4_Landscape of [CT]n,and [CTTT]n Microsatellite Density Peaks in the T2T-CHM13 assembly/S4.09.png]

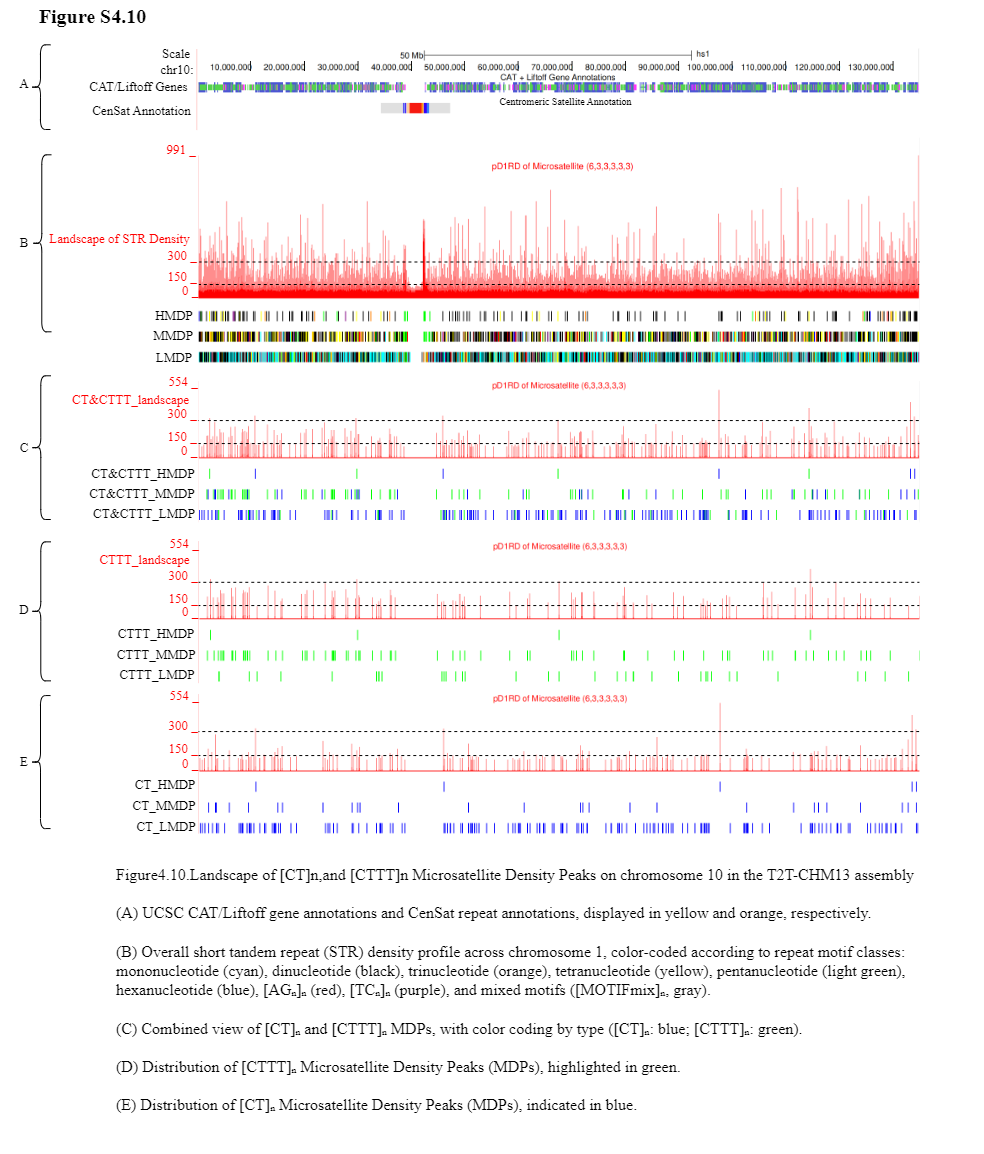

Supplement: Supplementary Table and Figure [file 713381_file03.zip › Fig and Table of Manuscript_20260322/Supplementary Figure/Supplementary Figure4_Landscape of [CT]n,and [CTTT]n Microsatellite Density Peaks in the T2T-CHM13 assembly/S4.10.png]

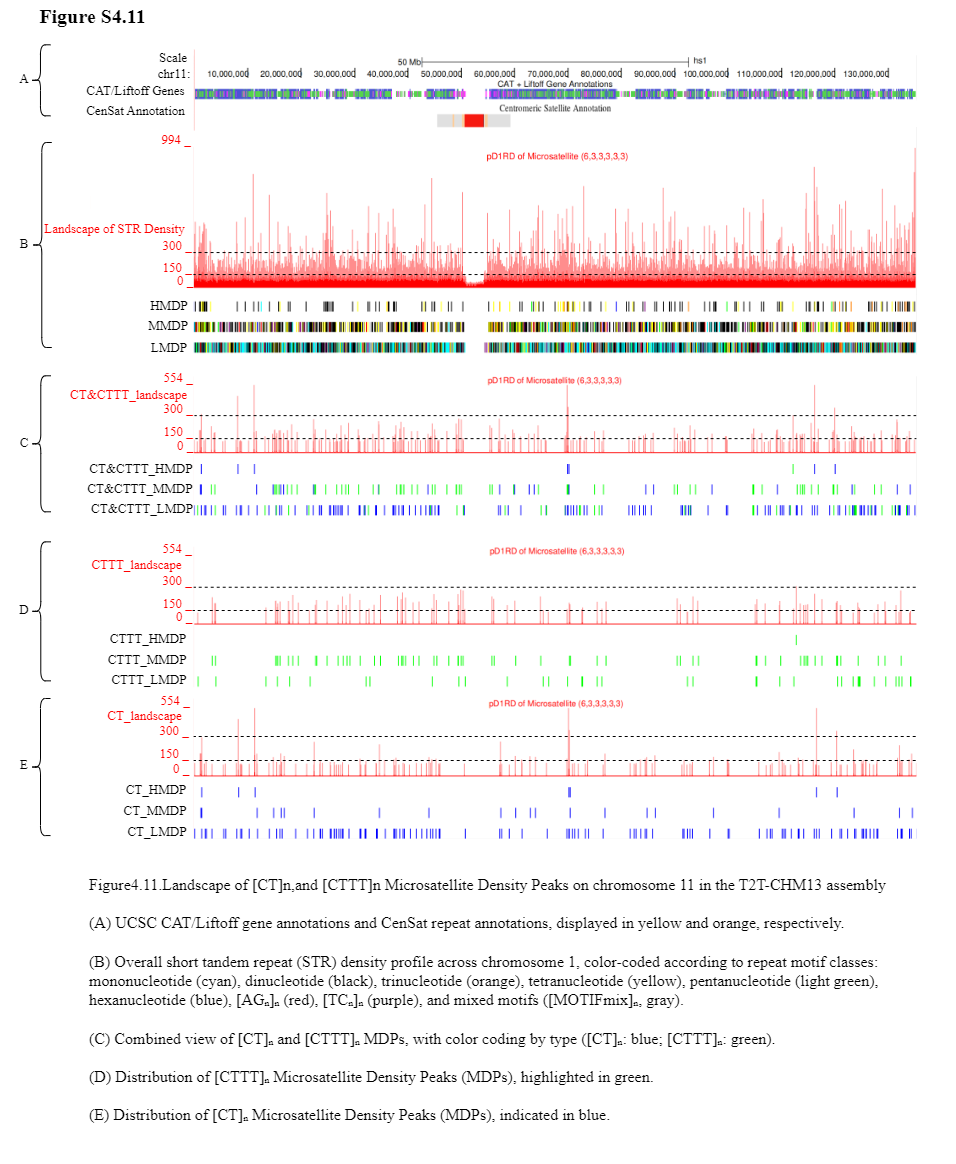

Supplement: Supplementary Table and Figure [file 713381_file03.zip › Fig and Table of Manuscript_20260322/Supplementary Figure/Supplementary Figure4_Landscape of [CT]n,and [CTTT]n Microsatellite Density Peaks in the T2T-CHM13 assembly/S4.11.png]

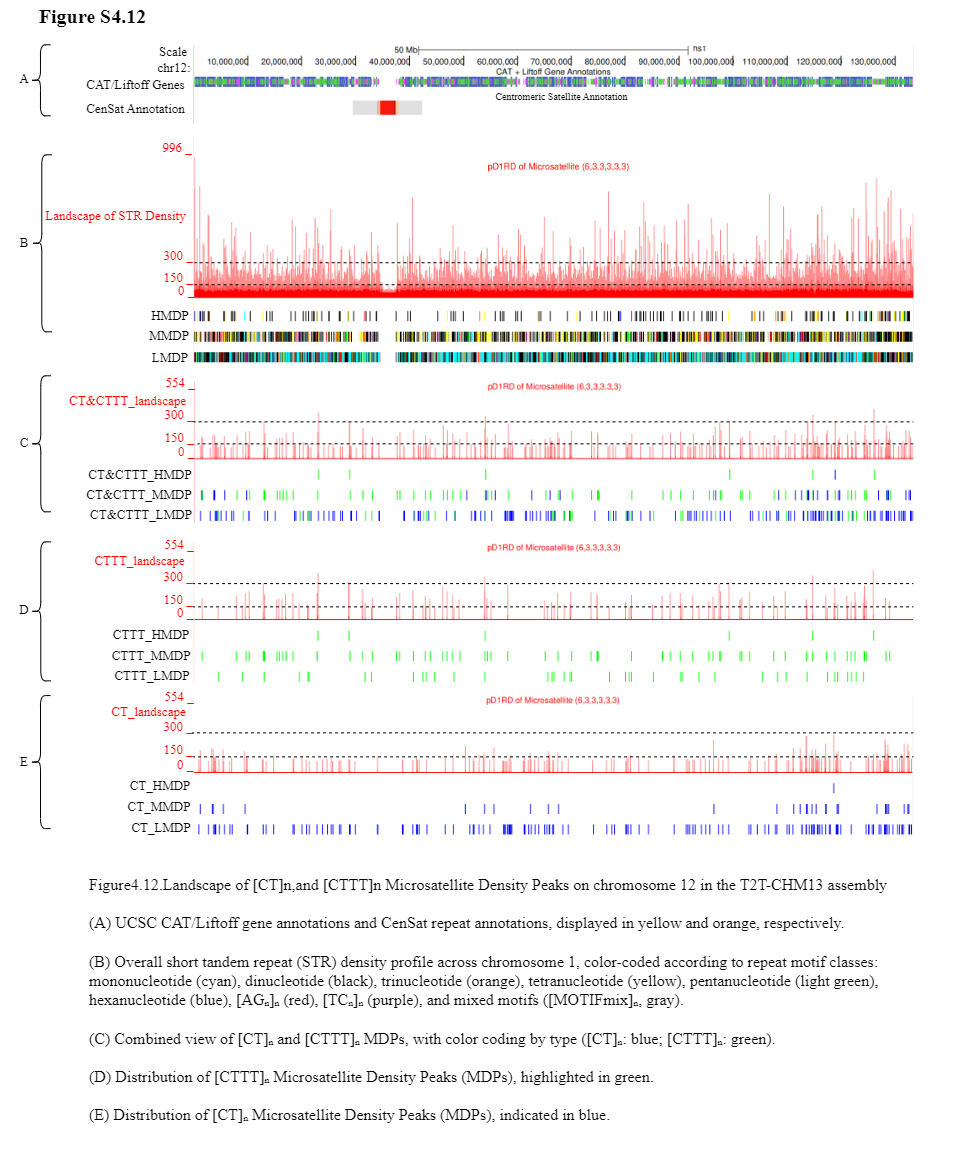

Supplement: Supplementary Table and Figure [file 713381_file03.zip › Fig and Table of Manuscript_20260322/Supplementary Figure/Supplementary Figure4_Landscape of [CT]n,and [CTTT]n Microsatellite Density Peaks in the T2T-CHM13 assembly/S4.12.png]

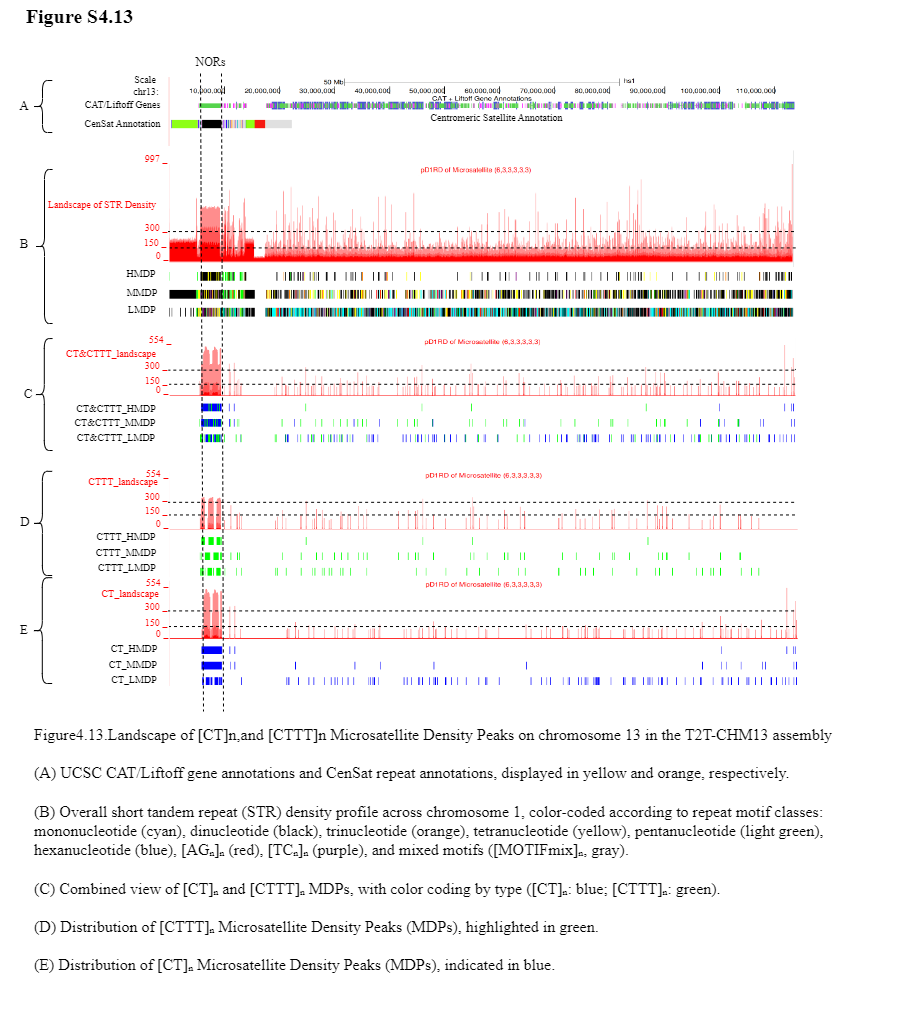

Supplement: Supplementary Table and Figure [file 713381_file03.zip › Fig and Table of Manuscript_20260322/Supplementary Figure/Supplementary Figure4_Landscape of [CT]n,and [CTTT]n Microsatellite Density Peaks in the T2T-CHM13 assembly/S4.13.png]

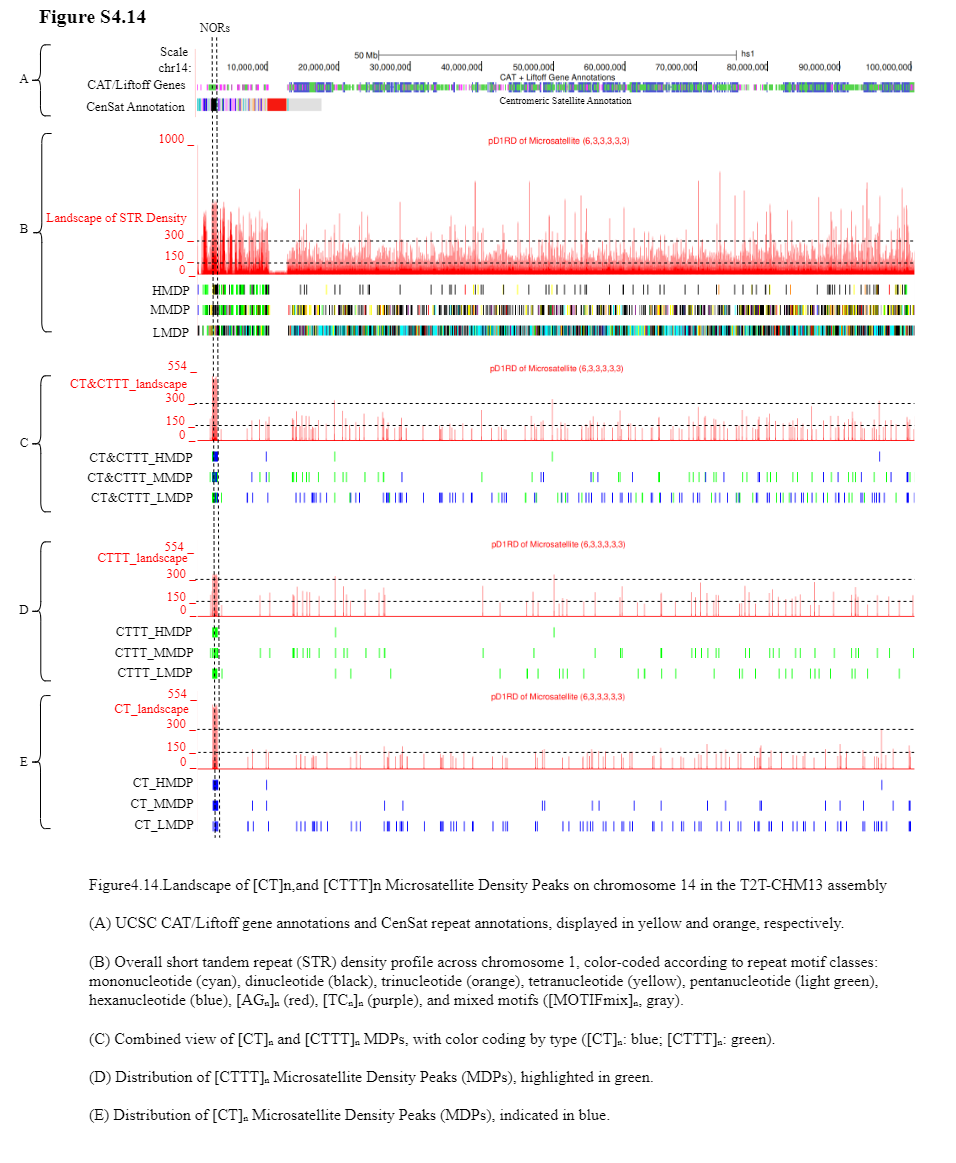

Supplement: Supplementary Table and Figure [file 713381_file03.zip › Fig and Table of Manuscript_20260322/Supplementary Figure/Supplementary Figure4_Landscape of [CT]n,and [CTTT]n Microsatellite Density Peaks in the T2T-CHM13 assembly/S4.14.png]

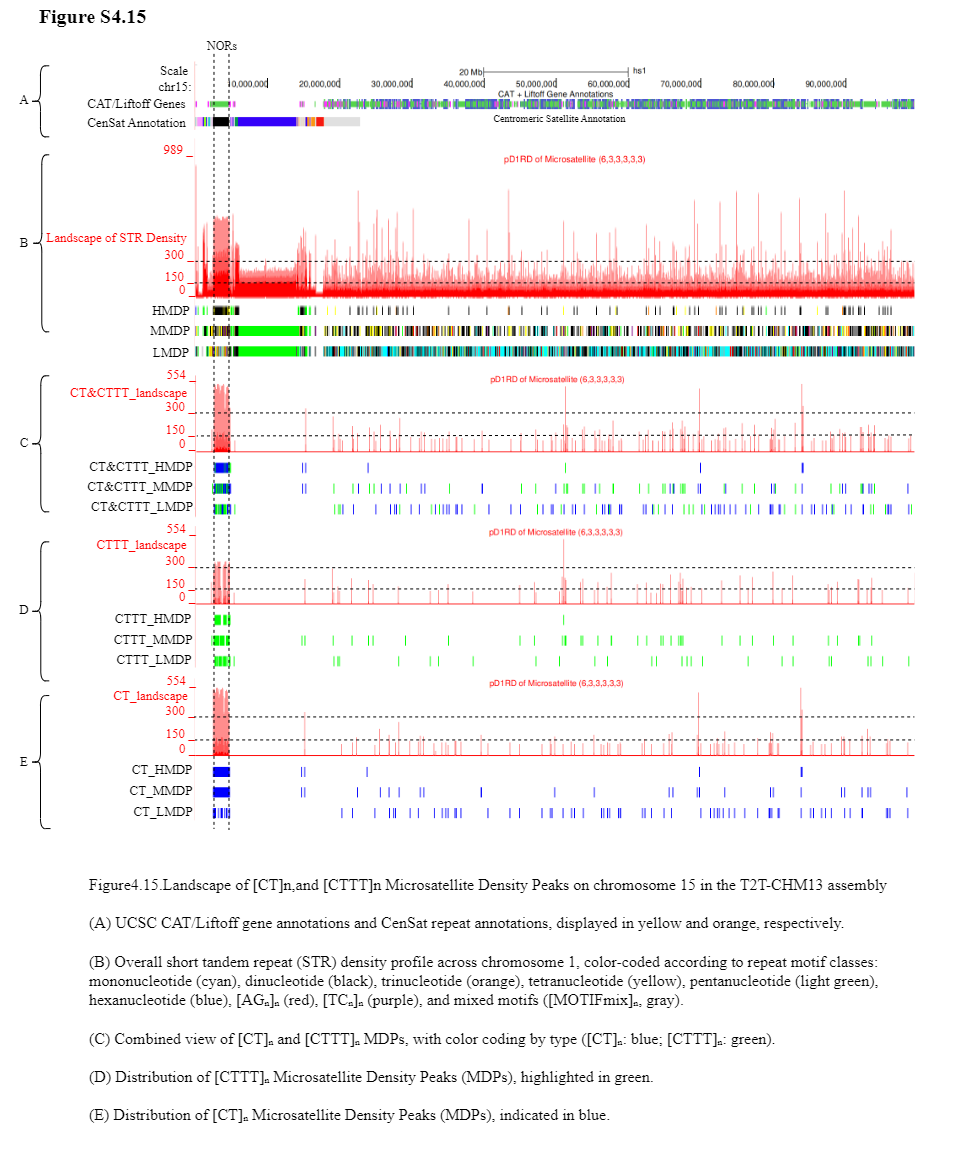

Supplement: Supplementary Table and Figure [file 713381_file03.zip › Fig and Table of Manuscript_20260322/Supplementary Figure/Supplementary Figure4_Landscape of [CT]n,and [CTTT]n Microsatellite Density Peaks in the T2T-CHM13 assembly/S4.15.png]

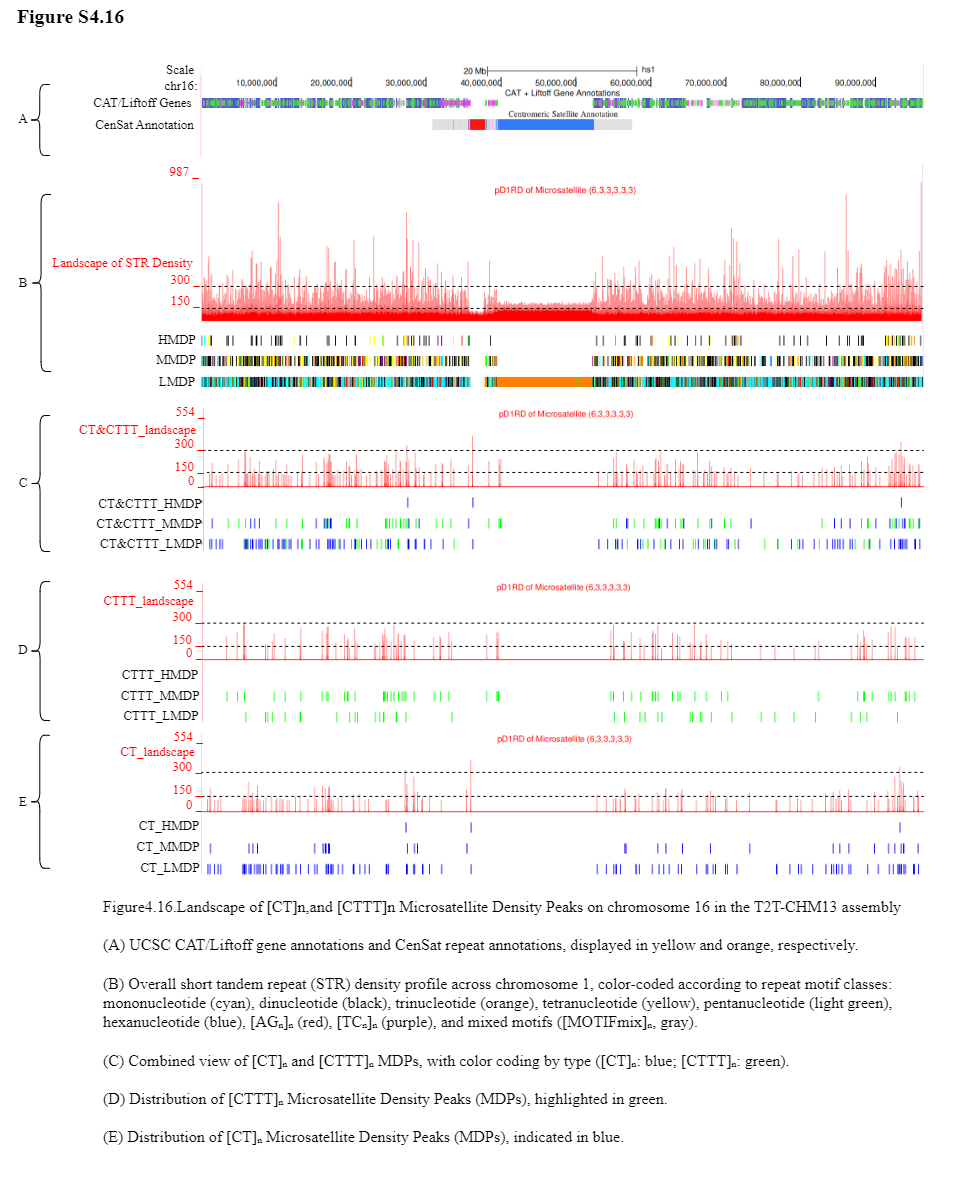

Supplement: Supplementary Table and Figure [file 713381_file03.zip › Fig and Table of Manuscript_20260322/Supplementary Figure/Supplementary Figure4_Landscape of [CT]n,and [CTTT]n Microsatellite Density Peaks in the T2T-CHM13 assembly/S4.16.png]

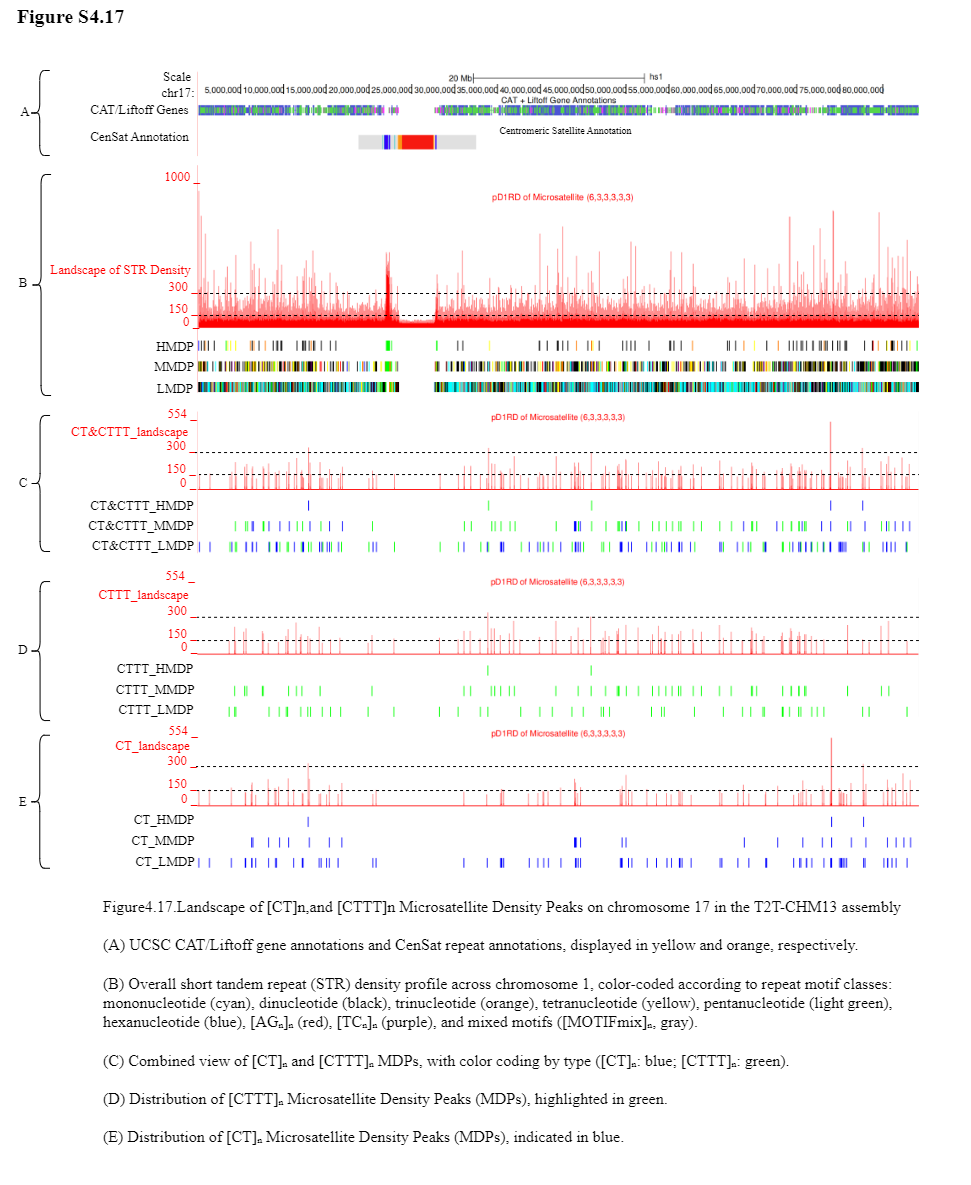

Supplement: Supplementary Table and Figure [file 713381_file03.zip › Fig and Table of Manuscript_20260322/Supplementary Figure/Supplementary Figure4_Landscape of [CT]n,and [CTTT]n Microsatellite Density Peaks in the T2T-CHM13 assembly/S4.17.png]

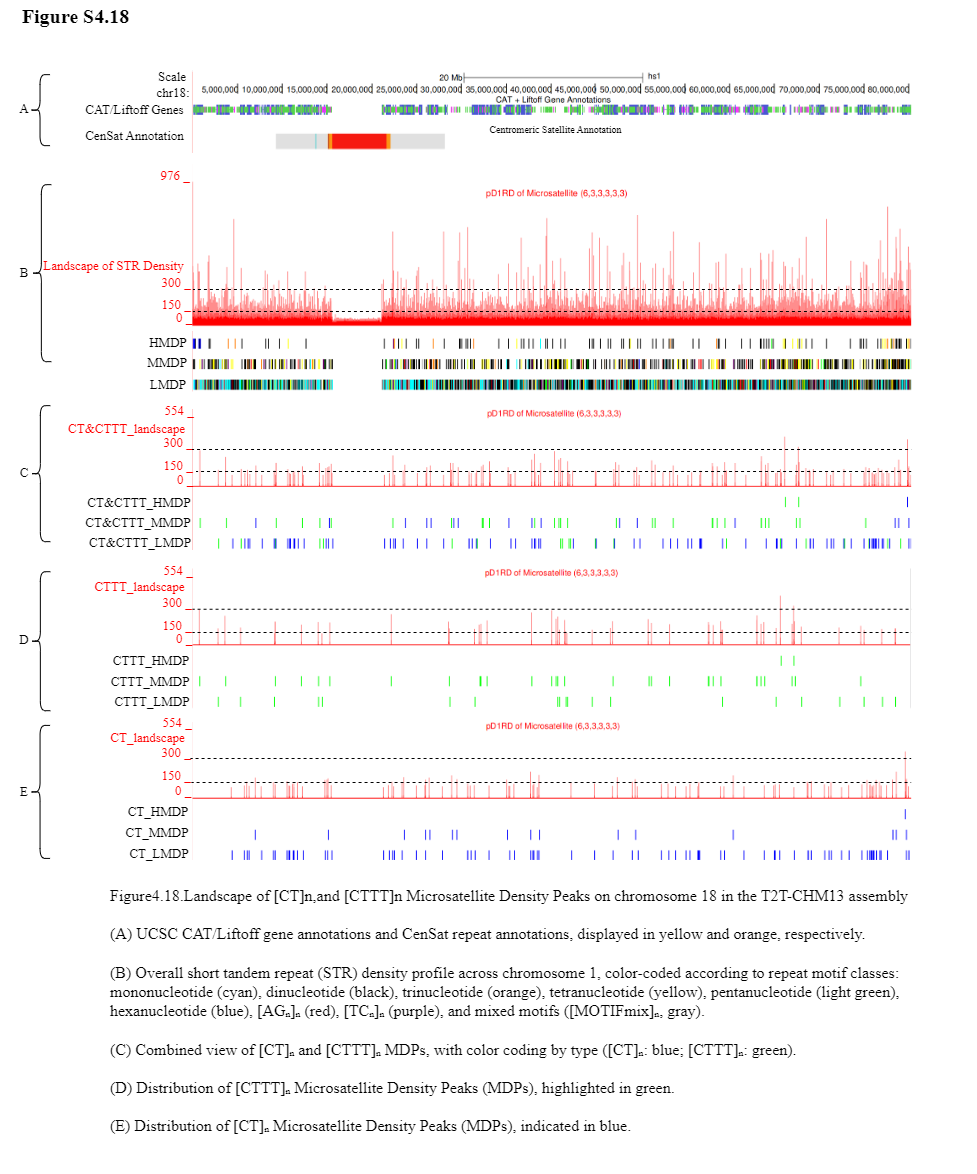

Supplement: Supplementary Table and Figure [file 713381_file03.zip › Fig and Table of Manuscript_20260322/Supplementary Figure/Supplementary Figure4_Landscape of [CT]n,and [CTTT]n Microsatellite Density Peaks in the T2T-CHM13 assembly/S4.18.png]

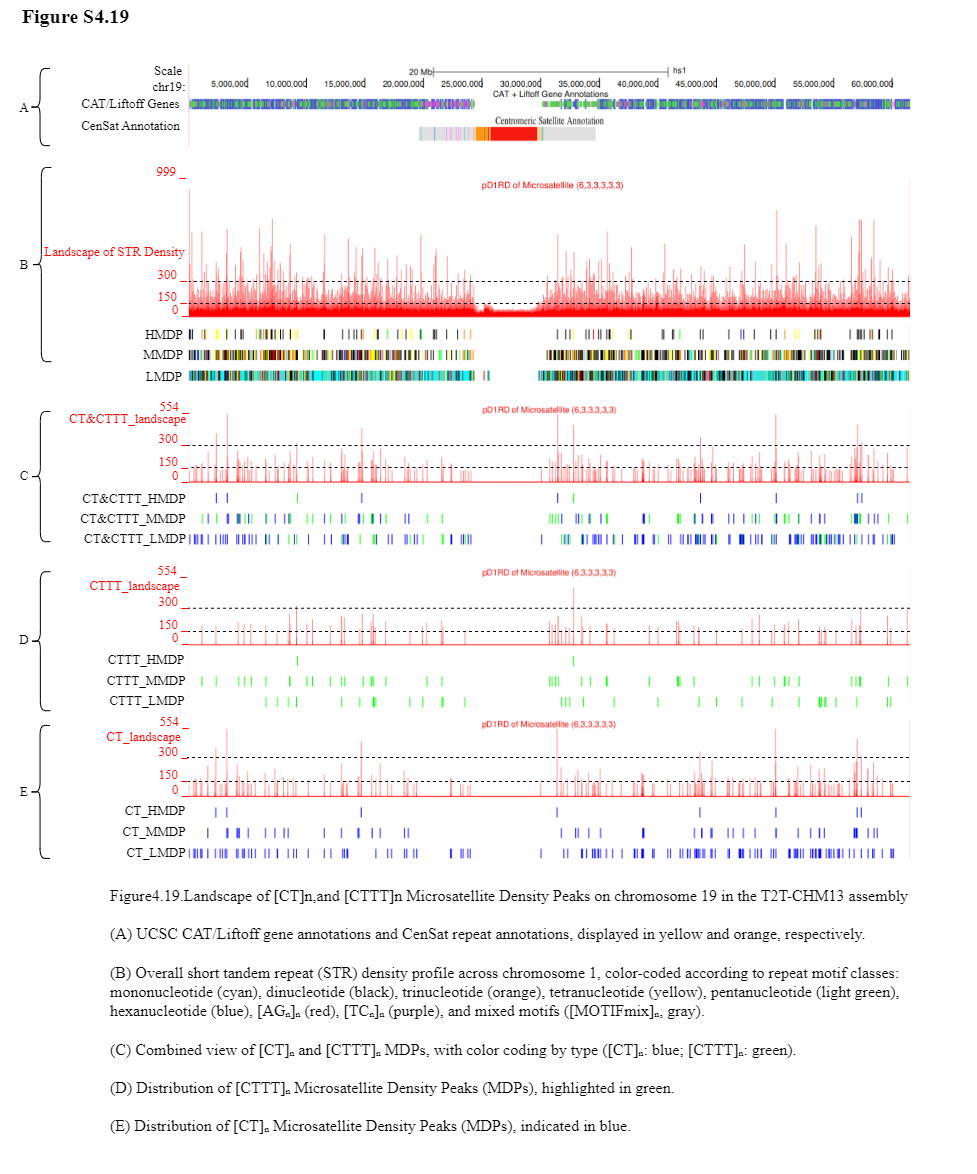

Supplement: Supplementary Table and Figure [file 713381_file03.zip › Fig and Table of Manuscript_20260322/Supplementary Figure/Supplementary Figure4_Landscape of [CT]n,and [CTTT]n Microsatellite Density Peaks in the T2T-CHM13 assembly/S4.19.png]

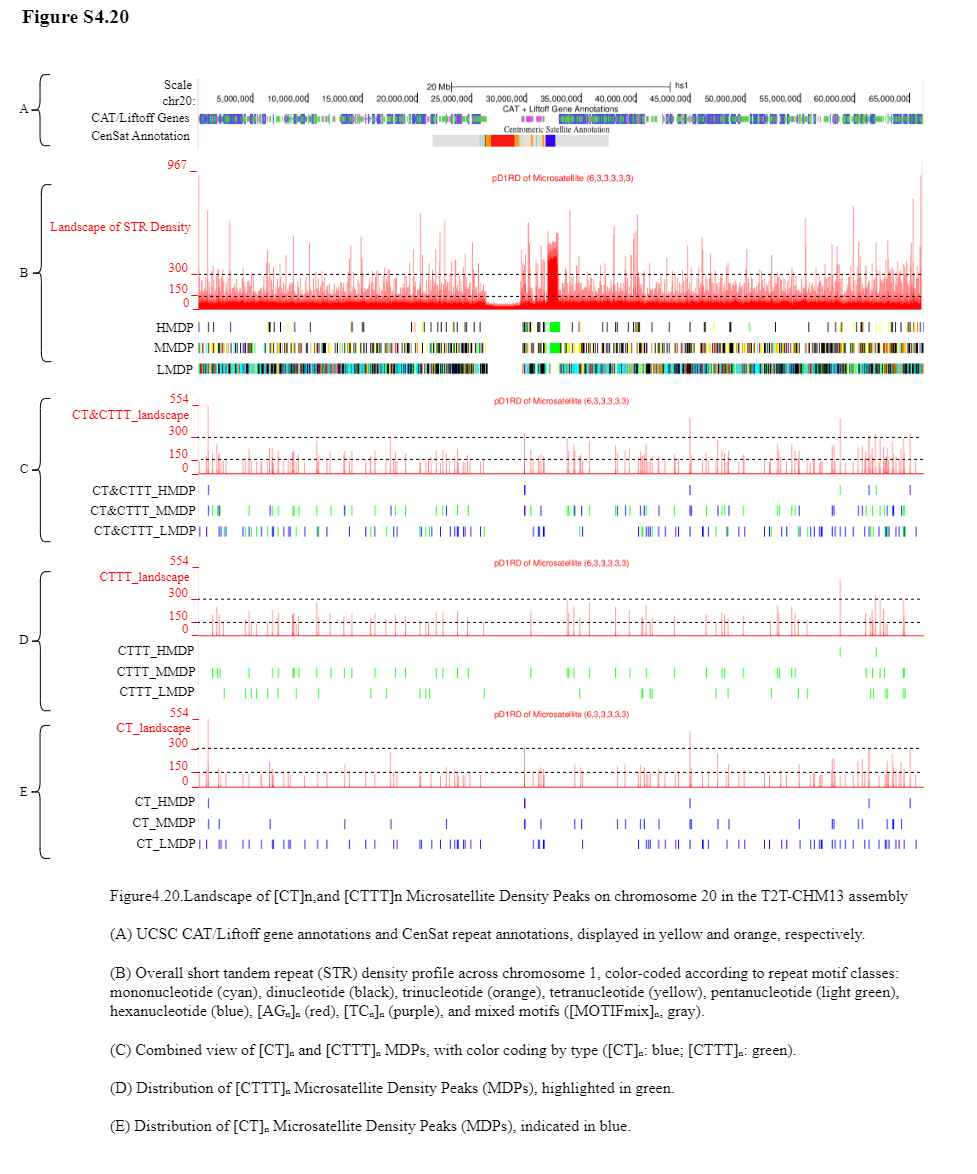

Supplement: Supplementary Table and Figure [file 713381_file03.zip › Fig and Table of Manuscript_20260322/Supplementary Figure/Supplementary Figure4_Landscape of [CT]n,and [CTTT]n Microsatellite Density Peaks in the T2T-CHM13 assembly/S4.20.png]

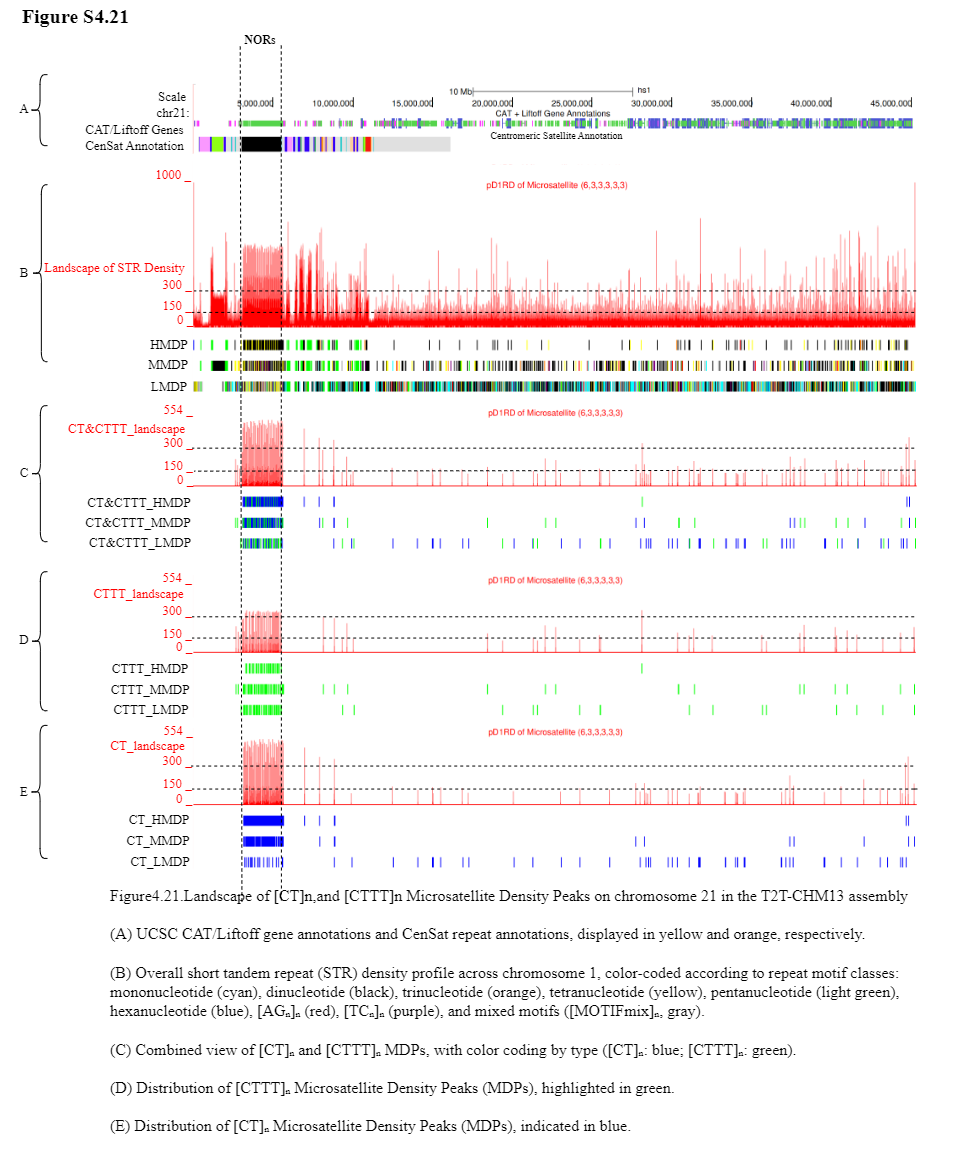

Supplement: Supplementary Table and Figure [file 713381_file03.zip › Fig and Table of Manuscript_20260322/Supplementary Figure/Supplementary Figure4_Landscape of [CT]n,and [CTTT]n Microsatellite Density Peaks in the T2T-CHM13 assembly/S4.21.png]

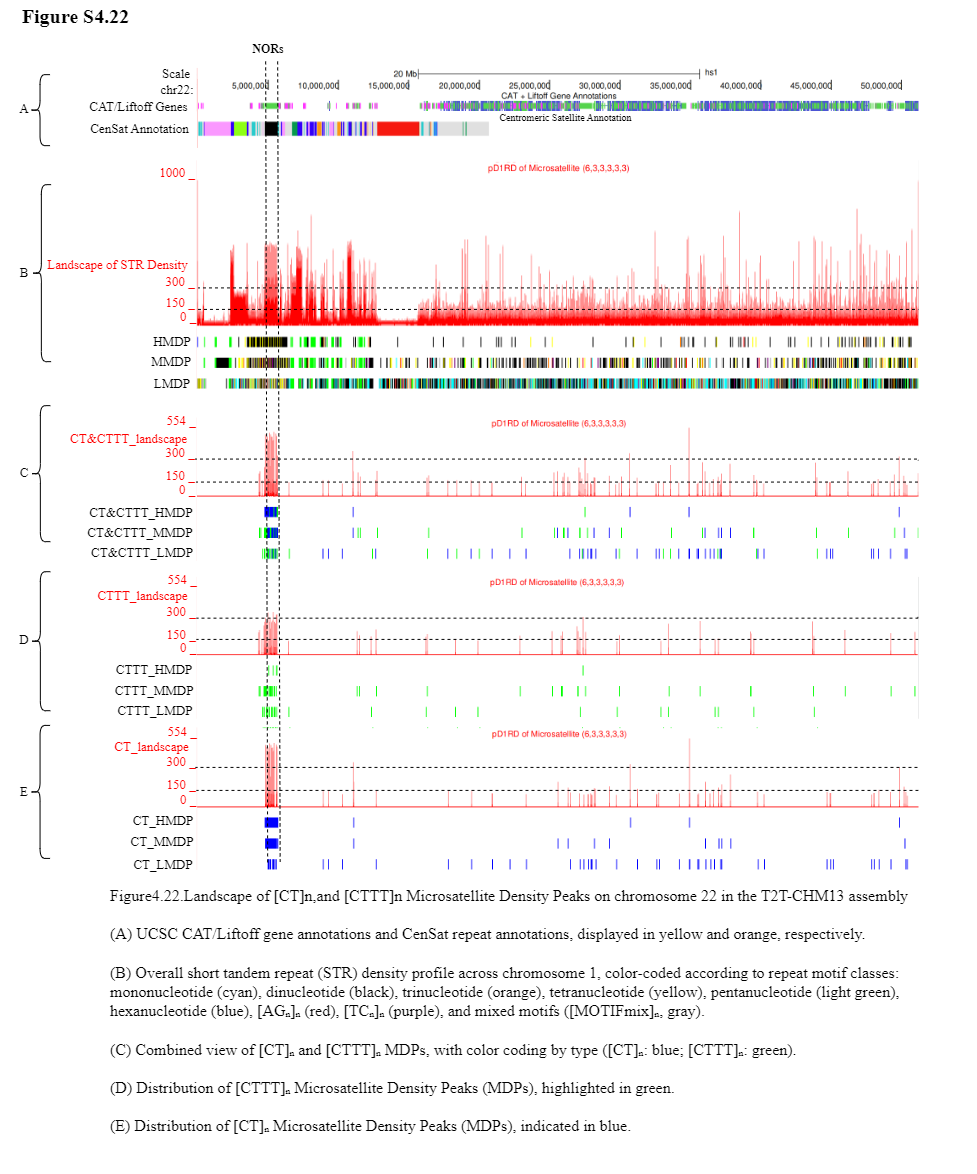

Supplement: Supplementary Table and Figure [file 713381_file03.zip › Fig and Table of Manuscript_20260322/Supplementary Figure/Supplementary Figure4_Landscape of [CT]n,and [CTTT]n Microsatellite Density Peaks in the T2T-CHM13 assembly/S4.22.png]

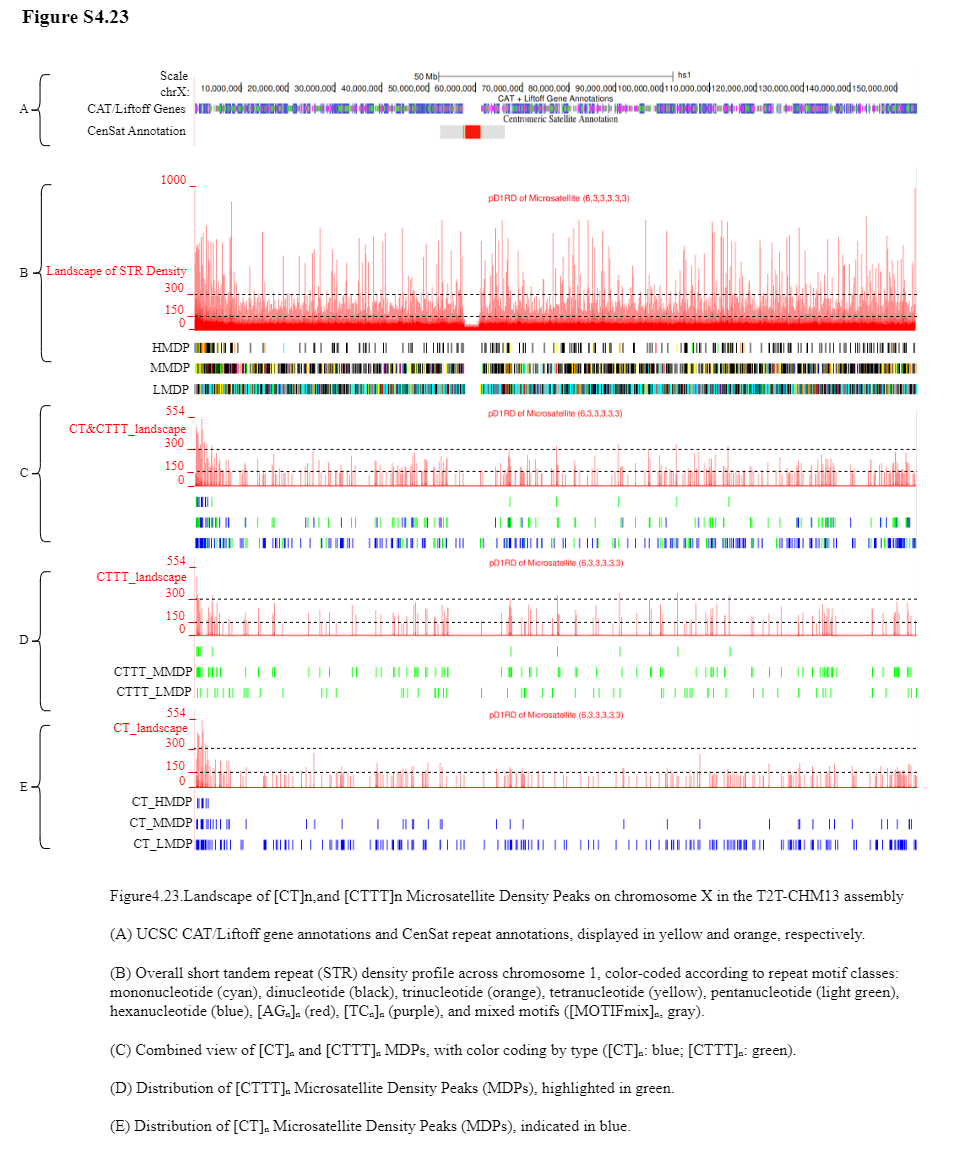

Supplement: Supplementary Table and Figure [file 713381_file03.zip › Fig and Table of Manuscript_20260322/Supplementary Figure/Supplementary Figure4_Landscape of [CT]n,and [CTTT]n Microsatellite Density Peaks in the T2T-CHM13 assembly/S4.23.png]

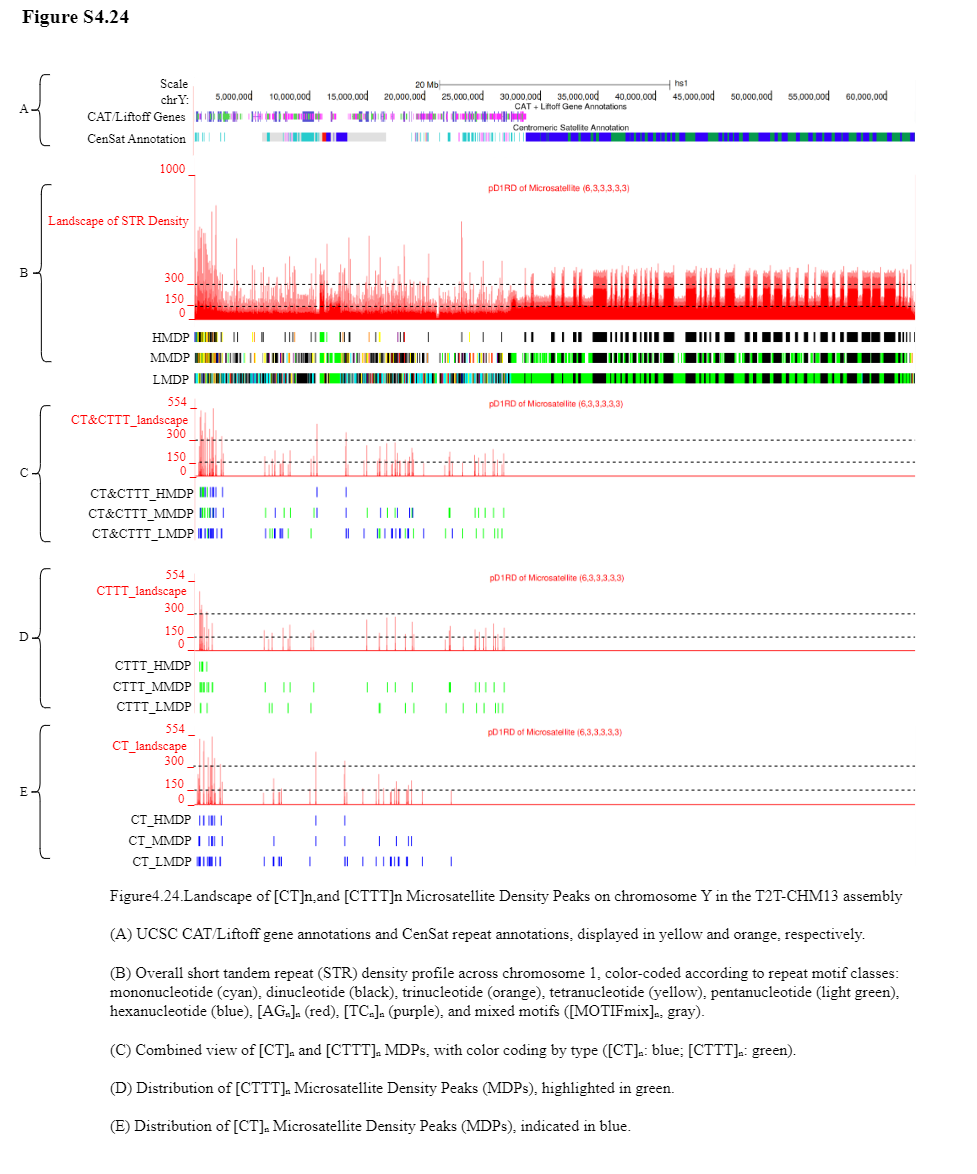

Supplement: Supplementary Table and Figure [file 713381_file03.zip › Fig and Table of Manuscript_20260322/Supplementary Figure/Supplementary Figure4_Landscape of [CT]n,and [CTTT]n Microsatellite Density Peaks in the T2T-CHM13 assembly/S4.24.png]
